# Coexistence of two divergent TprA/PhrA cell-cell communication systems in *Streptococcus mitis* coordinate bacteriocin production, competence, oxidative stress responses, and interspecies competition with *S. pneumoniae*

**DOI:** 10.64898/2025.12.10.693483

**Authors:** Bárbara Ferreira, Carina Valente, Ozcan Gazioglu, Hasan Yesilkaya, Shaw Camphire, N. Luisa Hiller, Raquel Sá-Leão

**Affiliations:** Laboratory of Molecular Microbiology of Human Pathogens, Instituto de Tecnologia Química e Biológica António Xavier, Universidade Nova de Lisboa (ITQB NOVA), Oeiras, Portugal; Polytechnic Institute of Castelo Branco, Castelo Branco, Portugal; School of Biological and Biomedical Sciences, University of Leicester, Leicester, UK; Department of Biological Sciences, Carnegie Mellon University, Pittsburgh, USA

**Keywords:** Mitis group streptococci, cell-cell communication, comparative genomics, bacteriocin, TprA/PhrA

## Abstract

Cell-cell communication (CCC) systems are key regulators of bacterial behaviors and adaptation. The human upper respiratory tract is co-colonized by commensal and pathogenic streptococci, but how CCC systems mediate their interactions remains unclear. Here, we investigated the TprA/PhrA quorum-sensing system, composed of the transcription factor TprA and its cognate signaling peptide PhrA, across *Streptococcus mitis* and *S. pneumoniae*. Comparative genomics showed that this system is broadly distributed across both species, with most strains sharing identical *phrA* alleles that enable interspecies signaling. In both species, the *tprA/phrA* module is commonly linked to the streptococcin E *(sce)* locus, encoding a putative bacteriocin. We show that activation of the *sce* operon enhances the competitive fitness of *S. mitis* in biofilm and infection models. In *S. mitis* strain C22, two diverse copies of *tprA/phrA* are present and differentially regulated, coordinating expression of *tprA/phrA* and the downstream *sce* locus through both shared and independent pathways. Transcriptomic analyses revealed redundancy, additivity, and cross-regulation between the two systems, linking them to the control of bacteriocin production, competence, and oxidative stress responses. Distinct promoter architectures and TprA-binding motifs underlie the functional divergence of these paralogues, highlighting how regulatory diversification can expand quorum-sensing outputs. Together, our findings show that *S. mitis* has evolved a flexible and layered communication network that integrates population sensing with antimicrobial and competence responses, providing a molecular basis for its competitive interactions with *S. pneumoniae* during colonization of the human respiratory tract.

## INTRODUCTION

The human upper respiratory tract consists of distinct regions with unique environments, each colonized by specific communities of bacteria, viruses, and fungi (1). Among common bacterial residents are streptococci, especially *Streptococcus mitis* and *S. oralis*, which establish early in life and persist as commensals (2). In immunocompromised individuals, these species can cause opportunistic infections (3). Akin to *S. mitis* and *S. oralis*, *S. pneumoniae* colonizes the upper respiratory tract of young children at high rates - up to 60% in Portugal, yet *S. pneumoniae* is typically cleared with age (4). In stark contrast to *S. mitis* and *S. oralis, S. pneumoniae* can also present as a major human pathogen (5). Dissemination from the upper respiratory tract to other tissues, such as lungs, middle ear, blood and brain, can cause mild to severe disease. Thus, the upper respiratory tract houses multiple related streptococci, with potential to interact with each other and potentially influence pathogenic outcome.

Recently, our group showed that certain *S. mitis* and *S. oralis* strains can inhibit and disrupt biofilms formed by diverse *S. pneumoniae* strains (6). This antagonistic activity is partly due to the production of antimicrobial peptides called bacteriocins, frequently regulated by CCC systems.

During CCC, bacteria release signaling molecules in response to environmental cues and population density. These signals coordinate group behaviors, enabling synchronized activation of genes involved in colonization, defense, and competition (7). In Gram-positive bacteria, CCC often relies on secreted peptides that either (i) trigger gene expression through two-component regulatory systems, or (ii) are imported via oligopeptide transporters and bind to transcriptional regulators (8, 9). The latter mechanism involves members of the RRNPPA family, consisting of the Rap, Rgg, NprR, PlcR, PrgX, and AimR protein families (8).

*S. pneumoniae* encodes multiple paralogues of Tpr/Phr, including the lineage-specific TprA2/PhrA2 and the broadly conserved TprA/PhrA (10, 11).In both cases, TprA is a transcriptional regulator and PhrA a signaling peptide that modulates TprA activity (12, 13). In characterized strains, the TprA/PhrA controls the expression of a putative lantibiotic locus with antimicrobial potential (12) (**Fig. 1**). Further, TprA/PhrA promotes both colonization and disease progression (13, 14). Our recent work revealed that the transporter PptAB and the protease Eep serve to temporarily coordinate the TprA/PhrA and Rgg/SHP pathways, suggesting global coordination of CCC systems during infection (10, 11).

**Figure 1.**
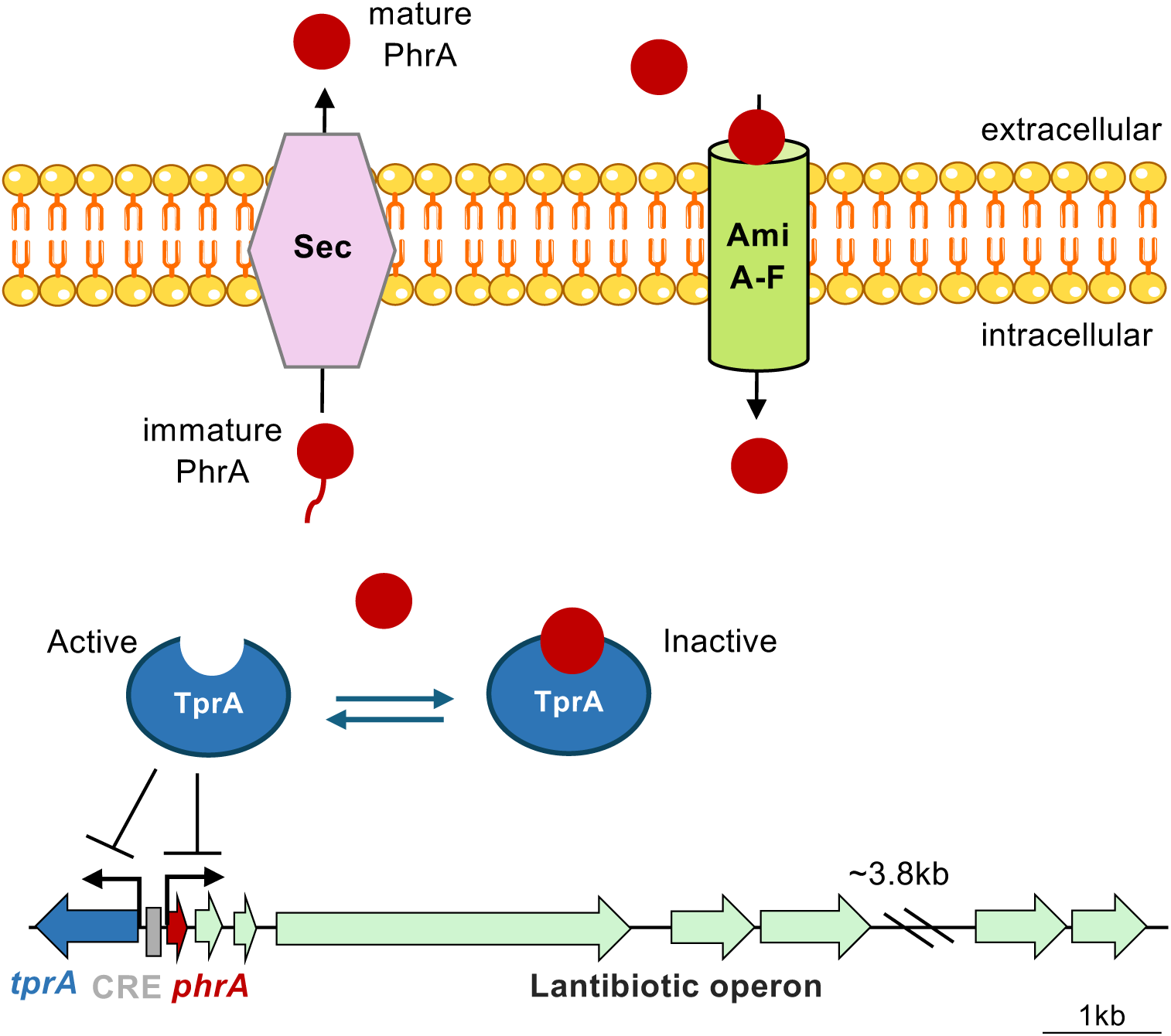
The *tprA/phrA* system controls gene expression in *S. pneumoniae* and is also present in other streptococcal species. This cell-cell communication system operates in a density-dependent manner and was previously characterized in *S. pneumoniae* strain D39 (12). At low cell density, TprA functions as a repressor by binding to a putative TprA-box, in the intergenic region between *tprA* and *phrA*, thereby inhibiting its own expression and the expression of downstream genes, including *phrA* and a lantibiotic locus with potential antimicrobial activity. *PhrA* is synthesized as a precursor peptide, which undergoes cleavage and is exported to the extracellular environment via the Sec pathway. As cell density increases, the mature *PhrA* peptides are reimported into the cell, where they bind to TprA, leading to the derepression of gene expression. The presence of a CRE element in the intergenic region between *tprA* and *phrA* is also regulated by glucose repression. Repression by TprA is shown with lines with a perpendicular bar. Genes and regulatory elements are color-coded: blue – *tprA*; grey – CRE binding-site; red – *phrA*; green – putative lantibiotic locus. The locus is drawn to scale; for simplicity, a 3.8 kb region is not shown.

The above-mentioned studies have focused on *S. pneumoniae*, such that the role of TprA/PhrA in non-pathogenic streptococci and interspecies communication remains unknown. Thus, the extent to which TprA/PhrA-mediated species interactions can influence bacterial behavior and virulence during colonization and infection remains to be addressed.

Here, we report the presence of the TprA/PhrA CCC system in non-pneumococcal streptococci, with a focus on its regulatory role in the commensal *S. mitis* and its function in interspecies communication. Using a comparative genomics approach, we found both species-specific and shared signaling cues and demonstrated that *S. mitis* can produce signaling molecules capable of activating the TprA/PhrA system in *S. pneumoniae*. We show that the gene cluster most frequently found downstream of *tprA* encodes a bacteriocin and promotes fitness in an *in vitro* biofilm model and in an *in vivo Galleria mellonella* infection model. Additionally, transcriptomic analyses revealed that the presence of two copies of the *tprA/phrA* genes in an *S. mitis* strain originated a new regulatory network that controls genes implicated in bacterial competition, virulence, natural competence, and metabolism.

## MATERIALS AND METHODS

### Bacterial strains, peptides and growth conditions

The bacterial strains and peptides used in this study are listed in **Table S1.** For all assays, strains were streaked from skim milk-tryptone-glucose-glycerine medium (STGG) stocks onto Tryptic Soy Agar with 5% sheep blood (BBL Stacker Plates, BD) and incubated overnight at 37°C in 5% CO₂. Pre-inocula were prepared in either C+Y_YB_ medium (for transformation assays,(15)) or chemically defined medium with galactose (CDMgal, (16)) for all other assays. Cultures were grown to an optical density at 600 nm (OD_600_) of 0.5 and then stored at -80°C in 15% glycerol.

### Comparative genomics analysis of the *tprA/phrA* system across streptococci

To identify homologues of *tprA* and *phrA*, across streptococcal species, we employed the tBLASTn tool with default parameters, using TprA and PhrA amino acid sequences as query, and a query coverage threshold of 90%, and an amino acid identity cutoff of 80%. Analyses were performed using streptococcal genome datasets available in the PubMLST database (last accessed in August 2024, detailed in **Table S2**). For *S. pneumoniae* (n=7,548), a subset of genomes from the PubMLST Pneumococcal Genome Library was used (17). For *S. mitis*, genomes were obtained from the species-specific genome library (n=322) within PubMLST. Genomes for all other streptococcal species were sourced from the Oral Streptococci Genome Library (n=1937) from PubMLST.

To characterize allelic variation in *tprA* among oral streptococci, we extracted *tprA* coding sequences from our comparative genomics dataset. Within each species, a distinct variant was defined as any sequence differing by ≥1 amino-acid residue. We identified the following variants: *S. pneumoniae* (n=202), *S. mitis* (n=98), *S. pseudopneumoniae* (n=41), *S. oralis* (n=2), *S. parasanguinis* (n=10), and *S. suis* (n=11). Unique variants across species were aligned to derive a global consensus sequence, and for position-specific analyses all variants were subsequently realigned using this consensus as the reference. All alignments were performed in CLC Genomics Workbench using default parameters.

To search for the genetic clusters potentially regulated by the TprA/PhrA system, 20 kb regions in the opposite strand to *tprA* were extracted. When *tprA* was located on the positive strand, the sequences of interest were reverse complemented. Nucleotide sequences were annotated using PROKKA (18). Annotated genetic clusters were aligned, and their genetic content was compared. Phylogenetic analysis of these regions was performed with Qiagen CLC Genomics Workbench v9.5.1 (Qiagen). To infer putative functions, representative regions from each cluster were queried by BLASTn against the NCBI database. Candidate functions were further validated by reciprocal BLAST searches against the original regions. This approach led to the identification of downstream regions associated with (i) antimicrobial peptide loci, including the streptococcin E locus (*sce*) and lantibiotic-like (*lan-like*) regions; (ii) antimicrobial resistance within mobile genetic elements (AMR-MGE); (iii) ABC transporters with unspecific functions; and (iv) choline-binding proteins (CBPs). *Lan-like*, AMR-MGE, and ABC transporter regions were grouped according to gene presence/absence, applying thresholds of >90% query coverage and >80% nucleotide identity to define members of the same group. For the classification of CBPs, all identified protein sequences were aligned, and a phylogenetic tree was constructed using the neighbor-joining method with the Jukes–Cantor model. The analysis included the unique CBPs from the downstream regions of *tprA* (n = 44) together with reference proteins: PspC, CbpC, CbpD, CbpE, CbpK, CbpG, CbpI, CbpJ, CbpM, LytA, LytB, LytC, PcpA, and PspA from *S. pneumoniae* TIGR4 (accession number: NC_003028.3), CbpM from strain R6 (accession number: NC_003098.1), and Cbp1smi–14smi from *S. mitis* B6 (accession number: NC_013853.1).

To visualize the relationship between specific downstream gene clusters and strain relatedness on a species level, we annotated phylogenetic trees of *S. pneumoniae* and *S. mitis*. Each tree was produced from a pre-built PopPUNK database, defining strain relatedness by both core and accessory genes (https://genome.cshlp.org/content/29/2/304, https://www.bacpop.org/poppunk-databases/). The *S. mitis* PopPUNK database was built from the same PubMLST collection of 322 genomes as described above. The *S. pneumoniae* tree was built from over >40,000 genomes (https://www.pneumogen.net/gps/), including all 7,548 genomes described above, then trimmed to only display the 7,548 genomes analyzed in this study while maintaining phylogenetic branch length relationships between genomes.

The trees were visualized and annotated using iTOL. A custom Python script was used to produce TREE_COLORS iTOL annotation files which assigned a label and color to each genome’s node_ID based on presence/absence of TprA, and presence of a specific downstream gene cluster in each genome.

### Construction of genetically modified strains

Deletion of *tprA/phrA* copies was done by allelic replacement with an antibiotic-resistant gene flanked by lox66 and lox71 sites as previously described (6). Briefly, the flanking regions of the gene to be deleted were amplified, ligated by Gibson Assembly (NEB), and the fused amplicon was amplified with nested primers, gel-purified, and used for transformation.

For the construction of the reporter cassette P*phrA*:mNeonGreen-luc, the *phrA* promoter from *S. pneumoniae* D39 was PCR-amplified with BsaI restriction sites added to the 5′ ends of the primers. The amplified fragments were digested with BsaI and ligated into the corresponding site upstream of mNeonGreen in plasmid pJB11 (19). The ligation products were transformed into *E. coli* DH5α, and transformants were selected on LB agar with 4 μg/mL chloramphenicol. Colonies containing the correct plasmid were PCR verified. Plasmids were extracted, linearized with BamHI, and transformed into *S. pneumoniae* D39 wild type or D39Δ*phrA* strains. Integration of the vector occurred via homologous recombination into the *cil* locus. Final transformants were PCR verified. The primers used for the construction of the genetically modified strains are detailed in **Table S1.**

For transformation, bacterial pre-inocula were grown in C+Y_YB_ medium (pH 7.4) to an OD_600_ of 0.5, diluted 1:100 in the same medium, and grown until an OD_600nm_ of 0.1. At that point, 150 ng/mL of the amplicon of interest, and 300 nM of cognate CSP (GeneScript, USA, **Table S1**) were added, and cultures were incubated for 4h. For transformants selection, cultures were plated in blood agar plates containing 200 µg/mL of spectinomycin, 300 µg/mL of kanamycin or 8 µg/mL of chloramphenicol, according to its resistance profile. Colonies were PCR verified to evaluate if they carried the correct genetic manipulation.

### *In vitro* biofilm model

Single- and dual-strain biofilms were grown on an abiotic surface as previously described (16). Strains were grown in CDMgal until mid-exponential phase (OD600nm ∼ 0.5). A starting inoculum of 10^5^ CFU/mL for each strain was added to CDMgal supplemented with catalase (1600 U/mL). This mixture was then inoculated into 24-well culture-treated plates (Corning) and incubated at 34°C with 5% CO₂ for 24 or 48h. For dual-strain biofilms, strains were mixed in a 1:1 ratio, while single-strain biofilms were grown in parallel as controls using the same inoculum.

After incubation, biofilm supernatants were removed, and the biofilms were resuspended in sterile PBS. Serial dilutions were then plated on blood agar (BA) and BA containing either 300 µg/mL of kanamycin or 8 µg/mL of chloramphenicol to distinguish between strains in dual-strains biofilms. Plates were incubated at 37°C with 5% CO₂ atmosphere, and CFUs were counted the next day.

### In vivo Galleria mellonella competition infection model

Larvae were obtained from Livefood, UK, and those weighing between 25–30 mg with a white, milky appearance were selected for infection. To determine the lethal dose 50% (LD50), larvae were injected with *S. mitis* C22 wild-type (C22WT) strain at three bacterial loads, 1×10^5^, 5×10^5^, and 1×10^6^ CFU, in fresh PBS. Injections were performed using a 10 μL Hamilton syringe, administered into the last right proleg. The progression of infection and melanization was monitored. For the competitive infection assay, larvae were co-infected with a 1:1 mixture of C22 strains (WT versus Δ*sceABC* and Δ*phrA* versus Δ*sceABC*), each at an approximate inoculum of 5×10^5^ CFU in PBS. Infected larvae were placed in sterile Petri dishes and incubated at 37 °C with 5% CO2. At 6h post-infection (hpi), hemolymph was collected from 10 larvae per experimental group by gently squeezing the larvae. Each experiment was performed in quadruplicate. The hemolymph was serially diluted and plated on selective antibiotic agar to determine bacterial colony counts. The competitive index (CI) was calculated as the ratio of the two competing strains in the output CFU relative to their ratio in the initial inoculum.

### Cross-species signaling with luciferase reporters

*S. mitis* C22WT was grown in CDMgal medium and, at an OD_595_ of 0.3, the culture was divided into two: half was treated with 1 μM synthetic peptide PhrA1.1 C10 and the other half was left untreated. Both were incubated at 37°C for 30 min. Cultures were washed three time with PBS, growth was resumed for 1, 2 or 3h, and cell-free supernatants (CFS) were collected. CFS were added to reporter strains *S. pneumoniae* D39WT or D39Δ*phrA*, each carrying a transcriptional fusion of the D39 *phrA* promoter upstream of mNeonGreen and *luc* (P*phrA*:mNeonGreen-luc) integrated at the *cil* locus. For artificial induction of P*phrA*, synthetic PhrA1.1 C10 was added to a final concentration of 1 μM. Assays were performed in CDMgal medium containing 0.45 mg/mL luciferin (Abcam), and 1600 U/mL of catalase (Sigma-Aldrich) in 96-well plates at 37°C with 1s of orbital shaking. Bacterial growth (OD_595_) and luciferase activity (RLU/OD) were measured every 10min for 16h using a BioTek Neo2 Plate Reader. P*phrA* activation was indicated by the emission of light resulting from luciferin degradation upon transcription of the luciferase-encoding *luc* gene.

### RNA extraction and RT-qPCR

Strains were grown in CDMgal at 37 °C with 5% CO_2_ until reaching early exponential phase (OD_600nm_ ∼ 0.1). Cultures were then split into two groups: one treated with 1 μM synthetic peptide PhrA and the other left untreated (with DMSO) **(Table S1).** After a 2h incubation, cultures were immediately placed on ice, and RNA was extracted as described previously (20).

Absence of DNA contamination was confirmed by PCR targeting the *gapdh* gene using the extracted RNA as template. For RT-qPCR, mRNA was converted to cDNA using the cDNA Synthesis Kit (Bioline). These cDNAs were then used as templates in qRT-PCR with the SensiFAST SYBR Kit (Bioline), performed on a CFX96 Touch real-time PCR detection system (Bio-Rad).

Each experiment included three independent biological replicates, each with two technical replicates. The primers used for RT-qPCR are detailed in **Table S1.** Data analysis was conducted with Bio-Rad CFX software, and results were normalized using the housekeeping gene *gyrA*. Gene expression fold changes were calculated using the 2^−ΔΔCT^ method (21).

### RNA sequencing (RNA-seq)

Total RNA was extracted from three biological samples of strains C22 using growth conditions described in RNA extraction section (WT with and without stimulation by PhrA1.1 C10), C22Δ*tprA.I,* C22Δ*tprA.II*, and C22Δ*tprA.I*Δ*tprA.II.* Sequencing was conducted at Novogene using the Illumina NovaSeq6000 or NovaSeqX Plus platforms (paired-end, minimum 150 bp read length, 20 million reads per sample) and a prokaryotic directional mRNA library with rRNA removal.

RNA-seq data analysis followed a previously described workflow (22). Briefly, reads were mapped with Bowtie2 (23), converted from SAM to BAM format using Samtools (24), quantified with featureCounts (25), and analyzed for differential expression with the edgeR package in R. Genes were considered differentially expressed if they presented a false discovery rate (FDR) < 0.05, absolute log₂ fold change > 1.585, and P-value < 0.001. The data was further manually curated, with removal of genes with reads counts below 100, and inspection of fold changes at higher magnitude fold differences, for the characterization of a TprA genetic network.

The transcriptomic data from this study were submitted to the European Nucleotide Archive (ENA) under accession number PRJEB102718.

## RESULTS

### Orthologues of the TprA/PhrA signaling system are widely distributed across oral streptococci from mitis group

The TprA/PhrA signaling system described in *S. pneumoniae*, promotes colonization in the upper respiratory tract. Given this role, we investigated whether *tprA/phrA* is encoded in other oral streptococci and how it contributes to bacterial communication across diverse members of the upper respiratory tract microbiome.

Comparative genomics across curated genomes of oral streptococci representing species from the mitis, suis, anginosous, bovis, mutans, pyogenic, and salivarius groups **(Table S2)** revealed that orthologues of *tprA/phrA* are predominantly found in the mitis group, with low prevalence (<10%) in the suis group, and are absent in other streptococci groups **(Fig. 2A)**. Furthermore, within the mitis group, we found the highest prevalence of *tprA/phrA* homologues in *S. pneumoniae, S. pseudopneumoniae*, and *S. mitis*. Orthologues were also present in *S. parasanguinis*, *S. oralis*, and *S. gordonii* (**Fig. 2B**). In contrast, *S. australis* (n=2), *S. cristatus* (n=19), *S. infantis* (n=9), *S. rubneri* (n=2), and *S. sanguinis* (n=54) lacked *tprA/phrA* orthologues, although detection of orthologues in these species may have been hampered by the low number of genomes available. Notably, TprA sequences found in the mitis group streptococci revealed a high degree of amino acid conservation (**Fig. S1).**

**Figure 2.**
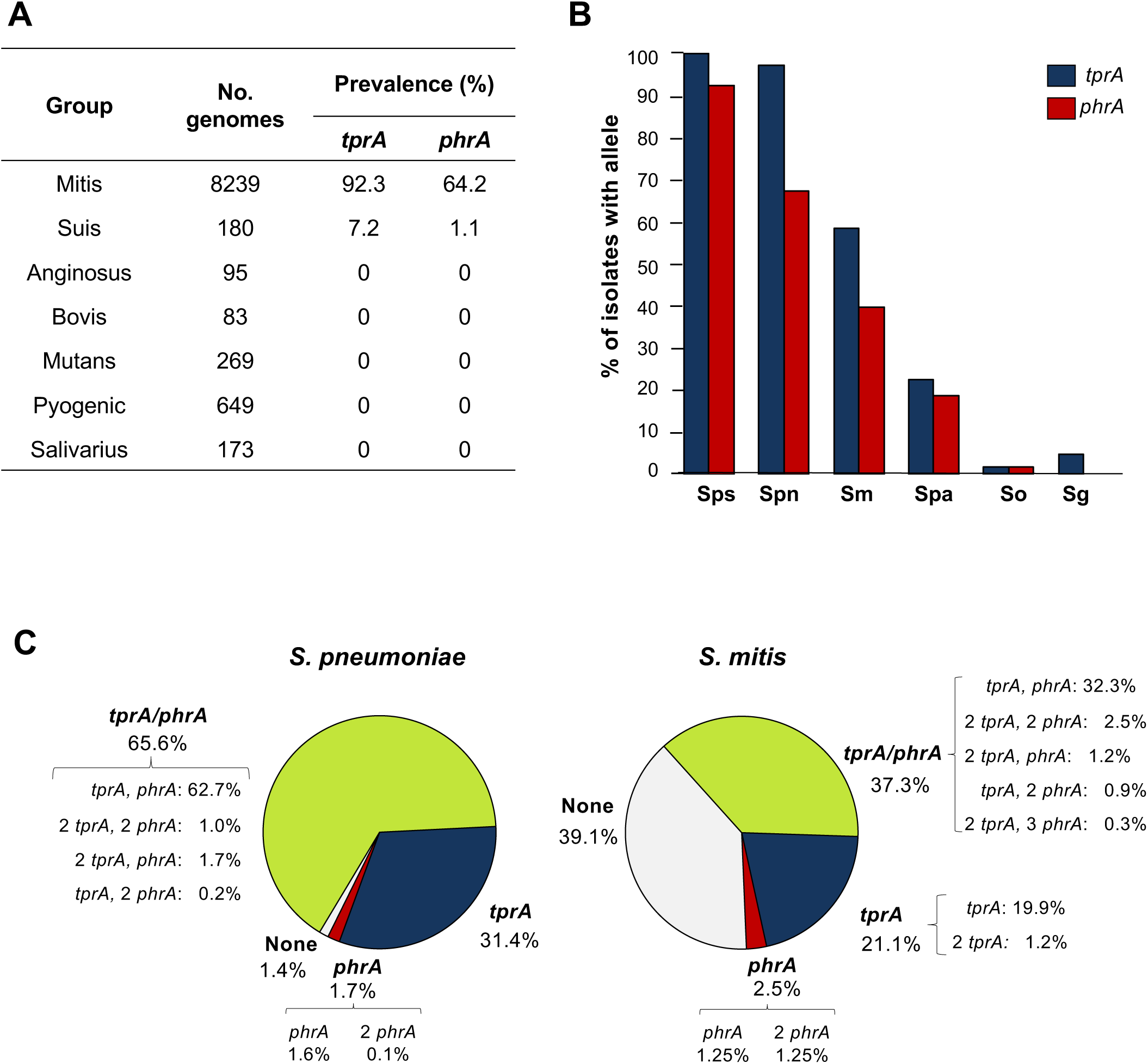
Distribution of the cell-cell communication system *tprA/phrA* among oral streptococci based on genomes publicly available at PubMLST. **A.** Prevalence of *tprA* and *phrA* homologues of D39 *S. pneumoniae* strain in oral streptococci. Tblastn cut-off values were 90% for query-cover and 80% amino acid identity. **B.** Relative abundance of *tprA* and *phrA* within streptococci of the mitis group. In parentheses the number of genomes screened for each species is indicated. In the x axis, from left to right: Sps - *S. pseudopneumoniae* (n=77), Spn - *S. pneumoniae* (n=7548), Sm - *S. mitis* (n=322), Spa - *S. parasanguinis* (n=53), So - *S. oralis* (n=108), and Sg - *S. gordonii* (n=41). **C.** Prevalence of *tprA* and *phrA* genes among 322 *S. mitis* and 7,548 *S. pneumoniae*.

As our main objective was to investigate interactions between *S. pneumoniae* and *S. mitis*, we focused on characterization of the TprA/PhrA system in these species. Interestingly, the genes encoding for the TprA regulator and its cognate PhrA peptide did not always co-occur. In *S. pneumoniae*, *tprA* and *phrA* alleles were present in 97.0% and 67.3% of the isolates, respectively, while in *S. mitis tprA* and *phrA* alleles were present in 58.4% and 39.8% of the isolates, respectively (**Fig. 2C).** The high proportion of genomes encoding only the receptor TprA suggests that PhrA is not always required for its regulation. In addition, 2.7% of *S. pneumoniae* and 5.2% of *S. mitis* genomes carried two copies of *tprA* raising the hypothesis that a second copy of the regulator could add another layer of genetic control, different from the one already described (13). Finally, 1.1% of *S. pneumoniae* and 4.6% of *S. mitis* genomes encoded two *phrA* alleles (and one genome encoded 3).

Altogether, these observations raised important questions about the evolution and functional significance of the TprA/PhrA system in both *S. mitis*, a commensal, and *S. pneumoniae*, a pathobiont.

### TprA/PhrA associated genes in *S. mitis* and *S. pneumoniae* are linked to bacterial warfare, host adhesion, antibiotic resistance genes, and ABC transporters

Previous studies on the TprA/PhrA regulatory system, particularly in pneumococcal strains D39 (model strain) and PN459-5 (PMEN1 lineage), established its role in regulating the downstream region which was identified as a putative lantibiotic locus (10, 12). To extend these observations, we analyzed the 20 kb downstream region of *tprA* in *S. mitis* and *S. pneumoniae* genomes in which there was no contig break in this region (i.e., 156 genomes in *S. mitis* and 5,952 genomes in *S. pneumoniae*, available at PubMLST).

The most prevalent genetic region downstream of *tprA* in both species was a streptococcin-associated locus, *sce* (**Fig. 3A, Tables S3-S4**). The longest version of this locus includes three genes: the bacteriocin gene *sceA*, its associated immunity protein *sceB*, and an ABC transporter *sceC* (26). Notably, in *S. mitis* and *S. pneumoniae*, most *sce* clusters lacked the bacteriocin gene *sceA* while retaining *sceBC* indicating most strains encode the immunity component, but only a few encode the bacteriocin. This pattern is consistent with a fitness cost for bacteriocin production, while maintaining immunity may provide a fitness advantage due to the high frequency of *sce* locus in both populations.

**Figure 3.**
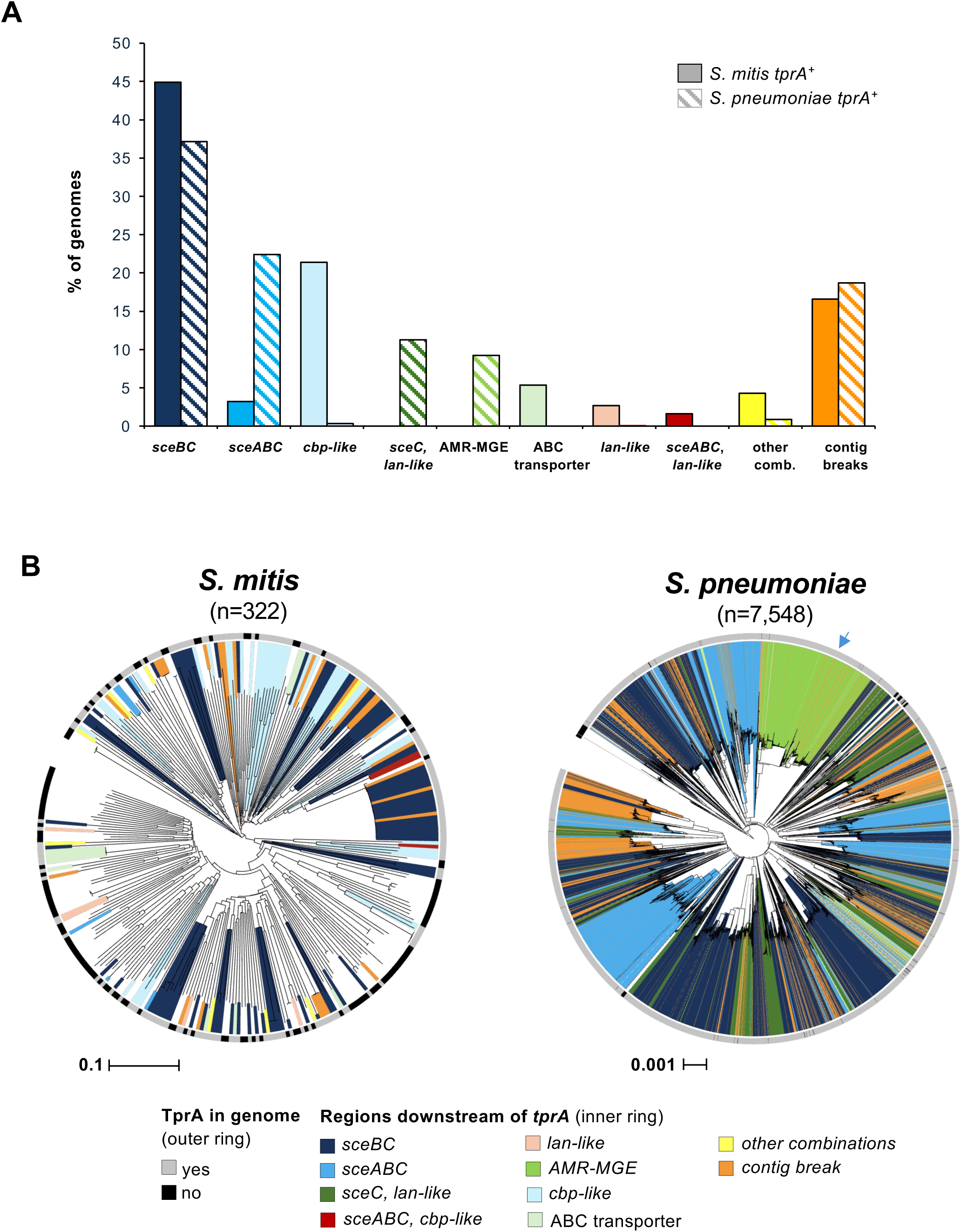
TprA/PhrA associated genes in *S. mitis* and *S. pneumoniae* are linked to bacterial warfare, host adhesion, antibiotic resistance genes, and ABC transporters. **A.** The most prevalent genetic regions downstream of *tprA.* These include streptococcin E (*sce*) *locus*, lantibiotic-like genes (*lan*-like), choline-binding proteins (*cbp*-like), antimicrobial resistance within mobile genetic elements (AMR-MGE) and ABC transporters’ encoding genes. Solid columns correspond to prevalence among *S. mitis* genomes that have *tprA* (187 *tprA^+^*); striped columns correspond to prevalence among *S. pneumonie* genomes that have *tprA* (7321 *tprA^+^*). **B.** Presence or absence of *tprA* and its downstream genetic content is shown for 322 *S. mitis* and 7,548 *S. pneumoniae* genomes. Outer rings indicate *tprA* presence (gray) or absence (black). The coloured inner rings represent the gene content within the 20 kb region downstream of *tprA*, grouped according to gene function. The arrow highlights evidence for horizontal gene transfer among the downstream region of *tprA*, with the presence of *cbp*-like region in an AMR-MGE (*tn916*-like) dominant clade. Trees were generated from Newick files from PopPUNK for *S. pneumoniae* and *S. mitis* and visualized via iTOL software.

Other types of loci were detected downstream of *tprA*, the most frequent ones encoding for choline-binding proteins (*cbp-like*, mostly present in *S. mitis*), lantibiotics (present in both species), and mobile genetic elements carrying antimicrobial resistance determinants (AMR-MGE, present in *S. pneumoniae*) (**Fig. 3A, Tables S3-S4**).

A more detailed analysis of the diversity of the choline-binding proteins identified downstream of *tprA* revealed nine distinct CBP categories in *S. mitis* (with *cbpJ-like* and *cbpGJ-like* being the most common) and four categories in the case of *S. pneumoniae* (with *cbpGJ-like* being the most common) (**Tables S3-S4, Fig. S2**). In pneumococci, CbpJ has been shown to be implicated with evasion to neutrophil-killing in lung infection setting (27), while CbpG can cleave fibronectin, that composes the host extracellular matrix (28). These proteins may therefore have an important role in both disease and colonization.

Three types of lantibiotic clusters were detected: types 1 and 2 were found only in *S. pneumoniae*, and type 3 was exclusive to *S. mitis* (**Tables S3-S4, Fig. S3)**. All lantibiotic clusters contained genes associated with lantibiotic bacteriocin production, including enzymes for modification and processing of the antimicrobial peptide, as well as genes for immunity and transport functions.

The AMR-MGE category was only found in *S. pneumoniae* and was classified into eight types (**Fig. S3)**. Types 1-7 belong to the Tn916 family. All regions within this category presented modules involved in conjugation, excision and integration of the mobile element in the bacterium genome, as well as genes associated with antibiotic resistance. Types 1, 2 and 6 contained genes encoding for resistance to macrolides, lincosamides and streptogramins (*ermB*) and tetracycline (*tetM*); type 3 contained *tetM* and *mefA* (associated with macrolide resistance), and types 4-5 and 7 contained *tetM* only. Type 8 category did not resemble the Tn916 family; instead, it contained genes encoding for a transposase and six ABC transporters with domains for antimicrobial resistance (e.g., ATP/binding cassette domain of the drug resistance transporter and related proteins, subfamily A) (**Fig. S3)**.

Two types of clusters classified as ABC transporters were detected in some *S. mitis*, yet no direct function could be attributed to them (**Fig. S3)**.

Finally, we overlaid the categories of genes found downstream of *tprA* on phylogenetic trees of *S. mitis* and *S. pneumoniae* generated based on their core genomes (**Fig. 3B**). In both species, the clusters of *sce* and lantibiotic genes are widely distributed across different lineages. Within specific clades of one species, we observed strains predominantly encoding the same gene cluster downstream *tprA* (for example, AMR-MGE cluster) and a few members of the same clade encoding a highly dissimilar gene cluster (for example, lan-like cluster) suggestive of horizontal gene transfer events. In *S. pneumoniae*, the Tn916-like element was limited to a single branch, demonstrating its point of entry into the population. Yet, there was also evidence of HGT with one strain within the Tn916-like containing also a *cbp-like* region.

In summary, the *sce* cluster, was the most frequent genetic locus downstream of *tprA* in *S. mitis* and *S. pneumoniae*. When combined with the putative lantibiotic locus, it suggests that one common function for *tprA/phrA* is regulation of bacterial warfare. Moreover, the presence of other loci suggests that this system may also regulate adhesion proteins, ABC transporters, and antibiotic resistance genes, highlighting its potentially broader impact on bacterial fitness and competition.

### The *sce* locus encodes a bacteriocin and requires PhrA

Considering the diversity of gene clusters downstream of *tprA*, and the fact that the *sce* locus was the most frequent among both *S. mitis* and *S. pneumoniae*, we investigated its potential antimicrobial effect in *S. mitis* using the recently described strain C22 (6). Strain C22 exhibits broad anti-pneumococcal activity, inhibiting more than 90% of a collection of 200 diverse pneumococcal strains (6), and carries a complete *sce* locus (**Fig. 4A**). Interestingly, strain C22 exhibits two copies of the *tprA/phrA* regulatory system, an occurrence identified in 2.5% of *S. mitis* isolates and 1.0% of *S. pneumoniae* strains **(Fig. 2C)**. In C22, one copy, here named *tprA.I/phrA1.1*, is located upstream of the full-length *sce* locus. The second copy, *tprA.II/phrA1.1\**, contains a *phrA1.1* allele with an early stop codon, and is encoded upstream of homologs to choline-binding proteins G and J **(Fig. 4A).**

**Figure 4.**
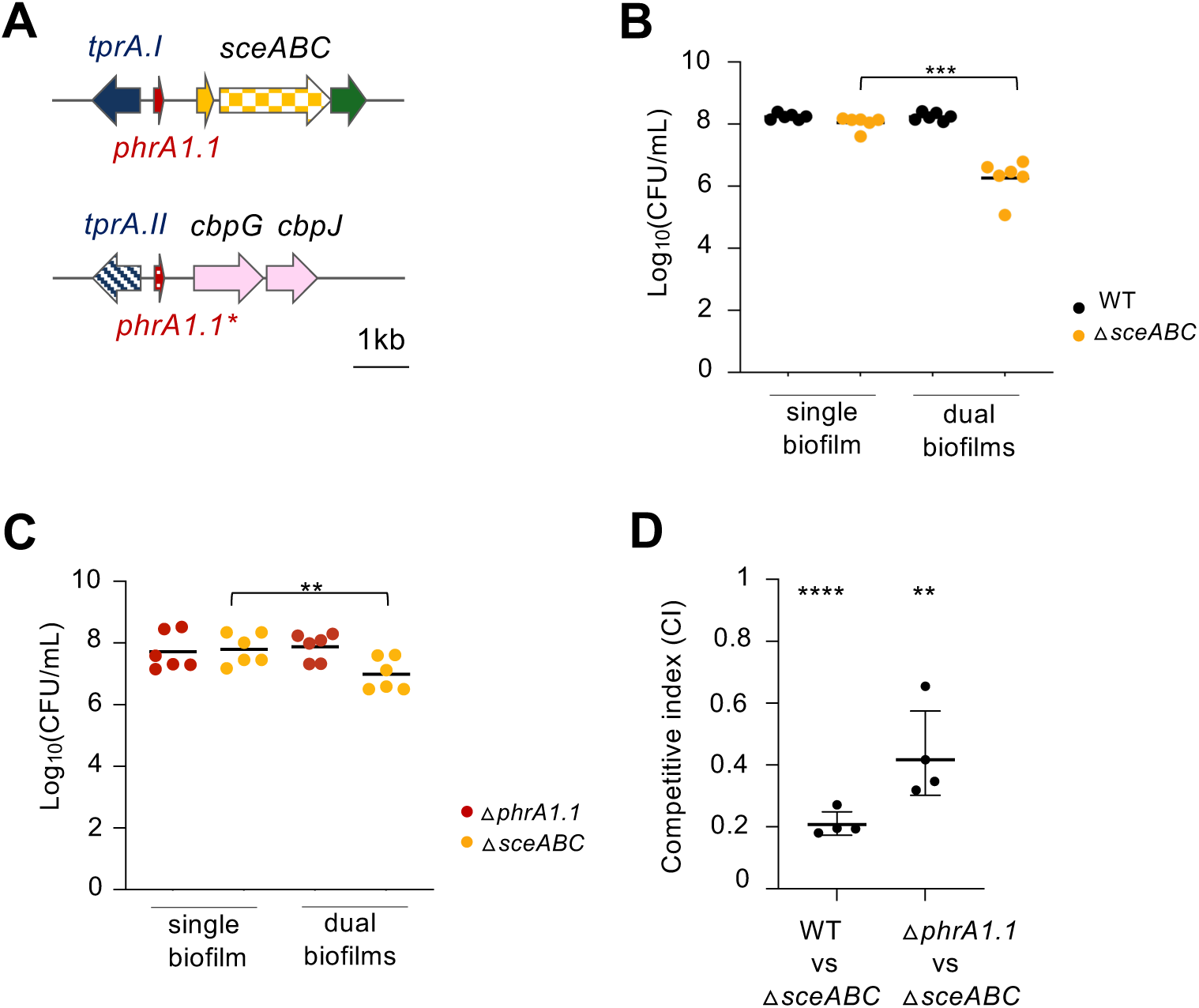
The *sce* locus exhibits antimicrobial activity, with PhrA regulating its inhibitory potential. **A.** Strain C22WT has two TprA/PhrA paralogues: one located upstream of the *sce* locus and other upstream of choline-binding genes CbpG and CbpJ. Genetic organization of both loci is shown. The *sce* locus comprises the bacteriocin gene *sceA*, its immunity gene *sceB*, and an ABC transporter gene *sceC*. Upstream, the *tprAI/phrA1.1* module regulates *sce* expression in response to the signaling peptide *PhrA1.1*. The other TprA/PhrA copy contains a non-functional *phrA1.1* allele due to an early stop codon. Arrows indicate gene orientation and transcriptional direction. **B.** The antimicrobial activity of the *sce* locus was assessed in an *in vitro* biofilm model by co-culturing C22WT and its isogenic △*sceABC* mutant at a 1:1 ratio (initial inoculum of 10^5^ CFU/mL) in CDM galactose for 48h. **C.** The role of PhrA on *sce* locus inhibition was evaluated using strain C22Δ*phrA (*Δ*phrA)* instead of WT. Bacterial loads quantification was done at 48h by plating the initial inoculum and the final biofilm on blood agar (total load) and blood agar supplemented with kanamycin (selective for △*sceABC* mutant). In panels B and C, P-values were calculated by paired Student’s t-test. Strains tested are color-coded: black – WT; yellow - △*sceABC*; red - △*phrA.* **D.** The *in vivo* role of Sce inhibition and its regulation by *phrA* was assessed using competition assays in the infection model *Galleria mellonella*. Competitive index (CI) values were calculated as the output ratio of mutant to wild-type bacterial load, normalized to the input ratio. Each dot represents pooled data from 10 larvae. P-values were calculated by one-sample Student’s t-test to a median hypothetical value of 1 (no fitness difference). \*\**P-value* <0.01; \*\*\**P-value* <0.0001. All assays were performed at least in triplicates, with graphics showing mean ± SEM.

The *tprA* paralogues in strain C22 exhibit over 90% amino acid conservation with each other - and with the *tprA* gene from the *S. pneumoniae* reference strain D39 - showing strong evolutionary conservation, as previously seen for TprA in both *S. mitis* and *S. pneumoniae* populations (**Fig. S1)**.

To evaluate *sce* locus inhibitory activity in a colonization-mimicking setting, we conducted *in vitro* biofilm competition assays. First, we competed the C22 wild-type strain (WT) against its isogenic *sce* deletion mutant (Δ*sceABC*, kanamycin resistant). While no significant inhibition of the Δ*sceABC* mutant strain was observed in 24-hour biofilms (**Fig. S4A**), after 48 hours the bacterial load of the Δ*sceABC* mutant was reduced by approximately 100-fold, confirming the inhibitory function of the *sce* locus (**Fig. 4B**).

Next, to assess whether the *sce* locus is regulated by PhrA, we performed biofilm competition assays using a *phrA* deletion mutant (Δp*hrA*) instead of WT, in competition with the Δ*sceABC* mutant. We hypothesised that the absence of the PhrA peptide pheromone would increase TprA-mediated repression of the *sce* locus in the Δ*phrA* mutant, thereby reducing expression of the streptococcin E locus and consequently decreasing competition against the Δ*sceABC* mutant. Indeed, this was observed in both 24- and 48-hour biofilms: inhibition of the Δ*sceABC* mutant strain was reduced by approximately 6-fold when co-cultured with C22Δ*phrA* compared to the levels observed with C22WT (**Fig. S4B, Fig. 4C**).

Thirdly, we investigated the role of *sce* locus inhibitory activity, as well as the ability of PhrA to induce this inhibition, in an infection setting. To this end, we performed the same competition assays in an *in vivo Galleria mellonella* infection model. The results showed that in co-infection experiments the Δ*sceABC* strain is outcompeted by the WT (CI=0.21, p-value < 0.0001); demonstrating that *sce* provides a competitive advantage in the model. Moreover, deletion of *phrA* buffered this effect (CI=0.43, p-value=0.0051), consistent with a role of PhrA in the regulation of *sce* (**Fig. 4D**).

Taken together, these findings demonstrate that *sce* is induced by PhrA, is involved in strain competition, and provides a fitness advantage in multi-strain infections.

### Streptococcal communication via TprA/PhrA is driven by shared and species-specific PhrA alleles

Having established that PhrA is a major contributor to the inhibitory activity of the *sce* locus in *S. mitis* strain C22, we next examined PhrA allelic diversity across *S. mitis* and *S. pneumoniae* populations to identify potential pherotypes. In *S. pneumoniae*, PhrA is processed during export, with the C-terminal 10 amino acids forming the active signaling peptide (11). Accordingly, we focused our analysis on sequence variation within this C-terminal region across both species.

Our analysis identified 16 PhrA alleles (**Fig. 5A, Tables S3-S4**). Among these, three variants (PhrA1.1, PhrA1.3, and PhrA1.6) were shared between *S. mitis* and *S. pneumoniae*. PhrA1.1 was the most prevalent variant in both species (**Fig. 5B**). PhrA1.1 regulates the lantibiotic loci in pneumococcal strain D39 (12); this same allele is present in strain C22WT.

**Figure 5.**
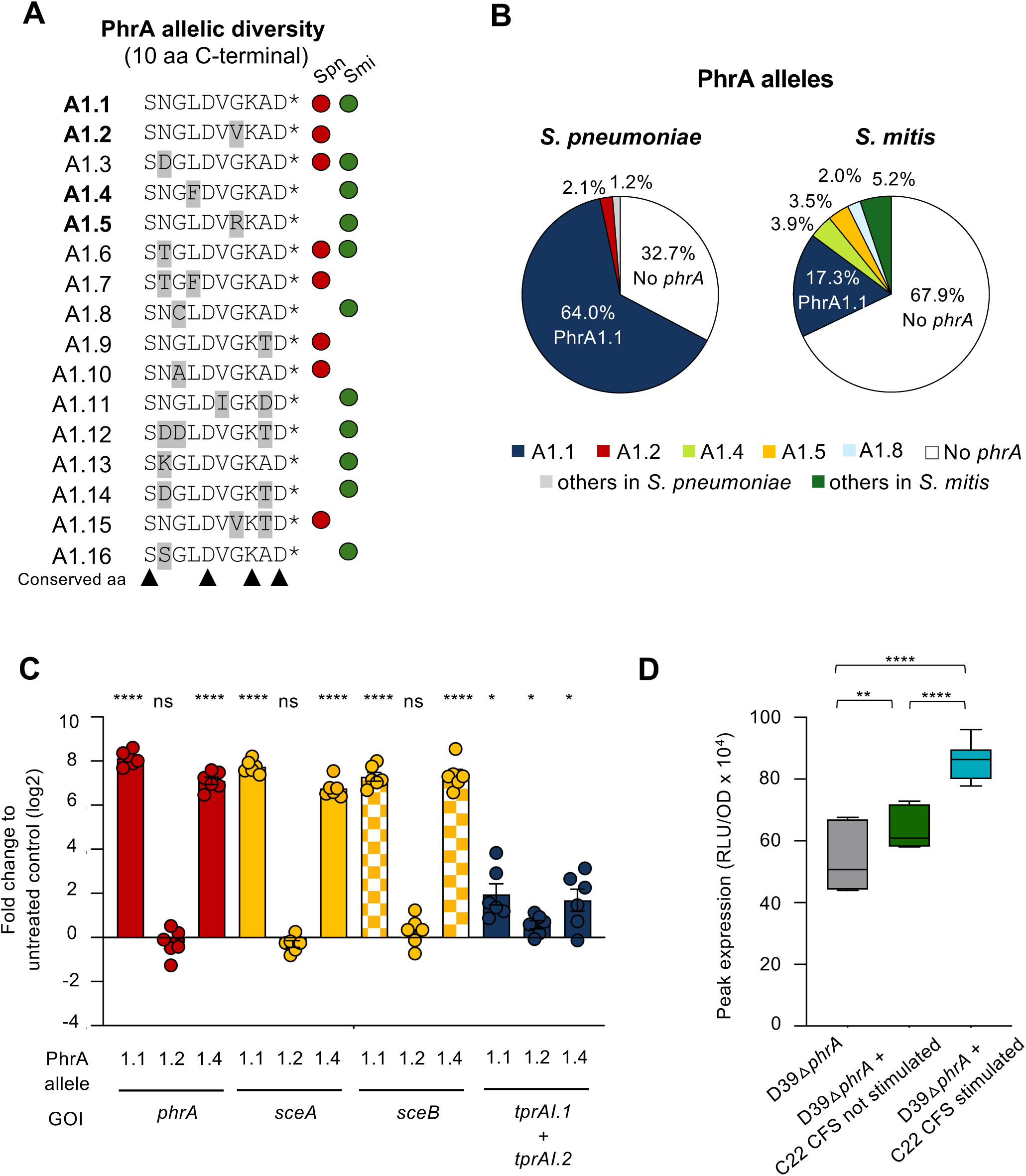
PhrA alleles drive streptococcal communication via the *tprA/phrA* system, involving both shared and species-specific variants. **A.** Diversity of PhrA mature peptides in 322 *S. mitis* and 7,548 *S. pneumoniae* genomes. Variants were defined based on amino acid differences in the last 10 residues. Differences from the PhrA1.1 variant (the most frequent) are highlighted in yellow. The positions of conserved amino acids are indicated by black triangles. Red and green circles indicate whether the allele was found in *S. pneumoniae* and/or *S. mitis* strains, respectively. **B.** Prevalence of PhrA allelic variants. Variants with >2% prevalence are shown individually, whereas those <2% were combined into the “other” category within each population. **C.** Assessment of PhrA allele strain specificity in *S. mitis* C22. The functionality of PhrA variants was evaluated by measuring induction of the *sce* locus using quantitative real-time PCR (qPCR). *S. mitis* C22WT cells were exposed for 30 min to synthetic C-10 PhrA peptides (PhrA1.1, PhrA1.2, PhrA1.4), and *sce* expression was quantified relative to untreated controls. **D.** Cross-species communication via the *tprA/phrA* system was evaluated using a bioluminescent reporter assay. *S. mitis* C22WT cultures were either pre-stimulated with 1 μM synthetic PhrA1.1 C10 or left untreated. Following treatment, cultures were washed to remove residual peptide and regrown for 3 hours before collecting cell-free supernatants (CFS). These supernatants were then used to assess activation of the *phrA* promoter in *S. pneumoniae* D39 Δ*phrA* reporter strain, carrying a PphrA:mNeonGreen-luciferase fusion at the *cil* locus. Reporter strains were incubated with either stimulated CFS, unstimulated CFS, or DMSO. Conditions tested are color-coded: grey (DMSO), green (3h CFS from non-stimulated C22WT), and cyan blue (3h CFS from stimulated C22WT). Data represent the mean ± SEM from three biological replicates. Data represent the mean ± SEM from three biological replicates. P-values were calculated by paired Student’s t-test. \**P-value* <0.05; \*\**P-value* < 0.01, \*\*\*\**P-value* < 0.0001; ns, not significant; GOI – gene of interest.

Analysis of PhrA alleles revealed distinct patterns of conservation and variability. The 1^st^, 5^th^, 8^th^, and 10^th^ residues were conserved across all variants, suggesting that these positions may be essential for PhrA interaction with its regulator, TprA. In contrast, residues at positions 2, 3, 4, 6, 7, and 9 were variable. Among these, three amino acid substitutions occurred at positions 2 and 3, two at positions 7 and 9, and one at positions 4 and 6. This pattern of variability implies that these sites are either less critical for TprA binding or may co-vary with polymorphic TprA alleles to maintain signaling compatibility (**Fig. 5A**).

To determine whether PhrA alleles confer strain specificity, we examined which PhrA variants are functional in C22 by assessing their ability to induce the *sce* locus. Using RT-qPCR, we measured *sce* expression in the absence of peptide and following the addition of synthetic C-10 PhrA peptides (**Fig. 5C**). Addition of PhrA1.1, the native allele of C22, resulted in a dramatic (>200-fold) increase in *sce* expression. Similarly, PhrA1.4, an allele specific to *S. mitis*, produced a comparable effect, indicating that *sce* expression in C22 can be triggered not only by its native allele but also by at least one additional allele encoded by other *S. mitis* strains. In contrast, PhrA1.2, which differs from both PhrA1.1 and PhrA1.4 at the seventh amino acid and occurs exclusively in *S. pneumoniae*, did not induce gene expression in C22, suggesting it is either nonfunctional or interacts specifically with a distinct TprA regulatory protein allele.

In the previous set of experiments induction of *tprA/phrA* gene expression was achieved with exogenous addition of PhrA peptides. To determine whether PhrA-mediated cross-species communication can occur directly between streptococci, we used an *in vitro* luciferase reporter assay targeting the *phrA* promoter (*PphrA*) in *S. pneumoniae* D39Δ*phrA* strains upon stimulation with cell-free supernatant (CFS) from *S. mitis*. As a control for the assay, we confirmed that both D39WT and the D39Δ*phrA* strains respond to synthetic PhrA peptide (**Fig. S5A**).

To achieve a high level of induction, we pre-stimulated C22 cultures with synthetic PhrA1.1 C10 before transferring them to PhrA-free medium for 1, 2, or 3 hours. CFS collected at these time points were then tested in the reporter assay (**Fig. 5D**, **Fig. S5B**). Supernatants from all time-points induced *PphrA* activity in D39Δ*phrA* compared to controls treated with unstimulated CFS or DMSO, demonstrating that signaling molecules released by *S. mitis* can activate *phrA* expression in *S. pneumoniae*.

Together, these results demonstrate that PhrA allelic diversity underlies shared species communication patterns among streptococci as the tested TprA receptors can respond to multiple PhrA variants, supporting inter-strain and inter-species signaling. Importantly, the ability of *S. mitis* supernatant to stimulate *S. pneumoniae* signaling confirms that PhrA functions as a diffusible cue enabling molecular dialogue between these species.

### The second copy of TprA is functional and responds to PhrA

The presence of two copies in strain C22 provided a unique opportunity to evaluate how this affected regulation of CCC systems. To determine whether the two *tprA* copies have distinct or overlapping roles in this regulatory pathway, we constructed deletion mutants for *tprA.I*, *tprA.II*, and a double mutant lacking both (τ<*tprA.I*τ<*tprA.II*). We then used RT-qPCR to quantify the expression of *tprA.*I, *phrA1.1,* and *sceA* and *secB* in the absence and presence of exogenously added PhrA1.1 (the functional phrA allele in strain C22). In the absence of exogenous PhrA, all deletion mutants showed increased *phrA.I* expression, indicating that both TprA copies act as inhibitors of *phrA.I* transcription. Deletion of *tprA.I* led to an 85- and 127-fold induction of *sceA* and *sceB*, respectively, while deletion of *tprA.II* caused 40- and 62-fold increases. The double mutant exhibited even stronger induction, with *sceA* and *sceB* upregulated 161- and 147-fold, respectively (p-values relative to Δ*tprA.I*: 0.0013 and 0.0910; relative to Δ*tprA.II*: 0.0069 and 0.0062) (**Fig. 6A**). Together, these results demonstrate that both *tprA* copies repress *sce* locus expression and exert additive regulatory effects.

**Figure 6.**
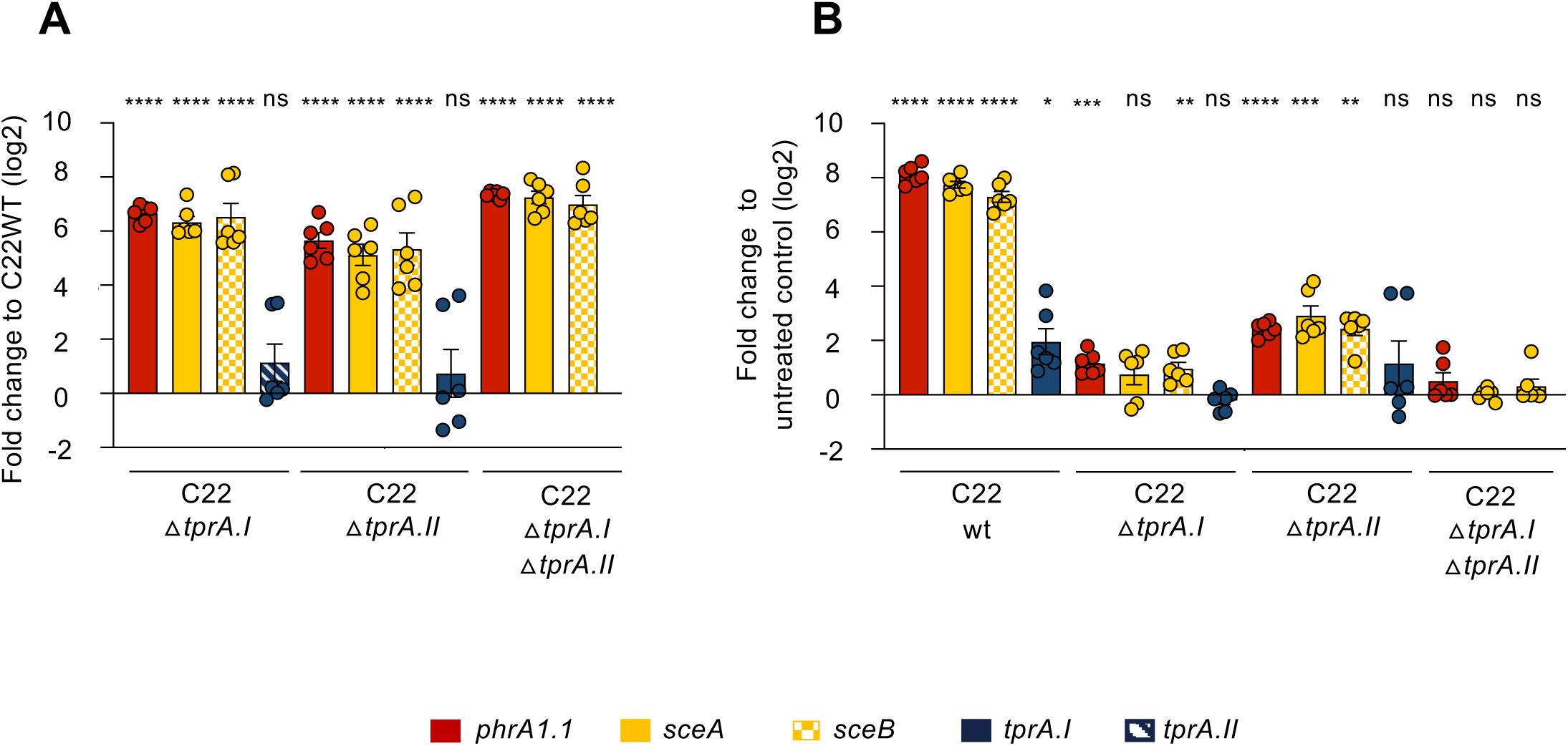
The presence of two copies of the TprA/PhrA system in *S. mitis* strain C22 modulates *sce* locus expression. **A.** The repressor role of TprA copies on *tprAI/phrA1.1.* and *sce* locus gene expression was evaluated in mutant strains C22Δ*tprAI,* Δ*tprAII*, and Δ*tprAI/tprAII* using qRT-PCR, with cultures grown in CDMgal media. The y-axis indicates the log₂ fold change in gene expression relative to the wild-type C22 strain. **B.** TprA responsiveness to exogenous PhrA1.1 was also tested and its impact on the expression of *tprAI/phrA1.1.* and *sce* locus evaluated. The y-axis indicates the log₂ fold change relative to the untreated control (no PhrA1.1 added). Genes and corresponding bars are color-coded in panels A-C as follows: solid blue – *tprA.I* transcription factor; stripped blue - *tprA.II* transcription factor, red - *phrA* pheromone; solid yellow - *sceA* bacteriocin; checkered yellow - *sceB* immunity protein; green - *sceC* ABC transporter; light pink - choline-binding proteins. Assays were performed in triplicate; graphs show means ± SEM, with two-way ANOVA, or paired t-test (\*\**P-value* < 0.01; **** *P-value* < 0.0001; ns, not significant).

We next repeated the same set of experiments in the presence of exogenously added PhrA1.1. The expression of *sceA* and *sceB* was higher in both the *ΔtprA.I* and *ΔtprA.II* backgrounds (>2-fold and >7-fold, respectively), when compared to untreated control conditions indicating that exogenous PhrA1.1 potentiates TprA.I and TprA.II derepression contributing to *sce* locus activation (**Fig. 6B**). Expectedly, in the double mutant, addition of this peptide had no influence on *sce* expression. Notably, exogenous addition of PhrA1.1 did not significantly affect expression of TprA.I nor TprA.II in the C22 deletion mutants.

We then hypothesized that, besides regulating downstream genes, the *tprA/phrA* system might also control expression of other genes. To identify the complete set of genes controlled by PhrA-mediated signaling, we performed RNA-seq and compared the transcriptomes for C22WT, with and without the addition of synthetic PhrA1.1 peptide. Consistent with our RT-qPCR results, *sceA*, *sceB*, *phrA1.1* and *tprA.I* were significantly upregulated with the addition of exogenous PhrA1.1. Notably, PhrA1.1 did not influence expression of *tprA.II* or *phrA1.1*,* suggesting different regulatory inputs for these paralogues. Finally, no other genes displayed differential expression after addition of PhrA1.1, suggesting PhrA-mediated signaling is limited to the downstream regulon in strain C22 (**Fig. 7A, Table S5)**.

**Figure 7.**
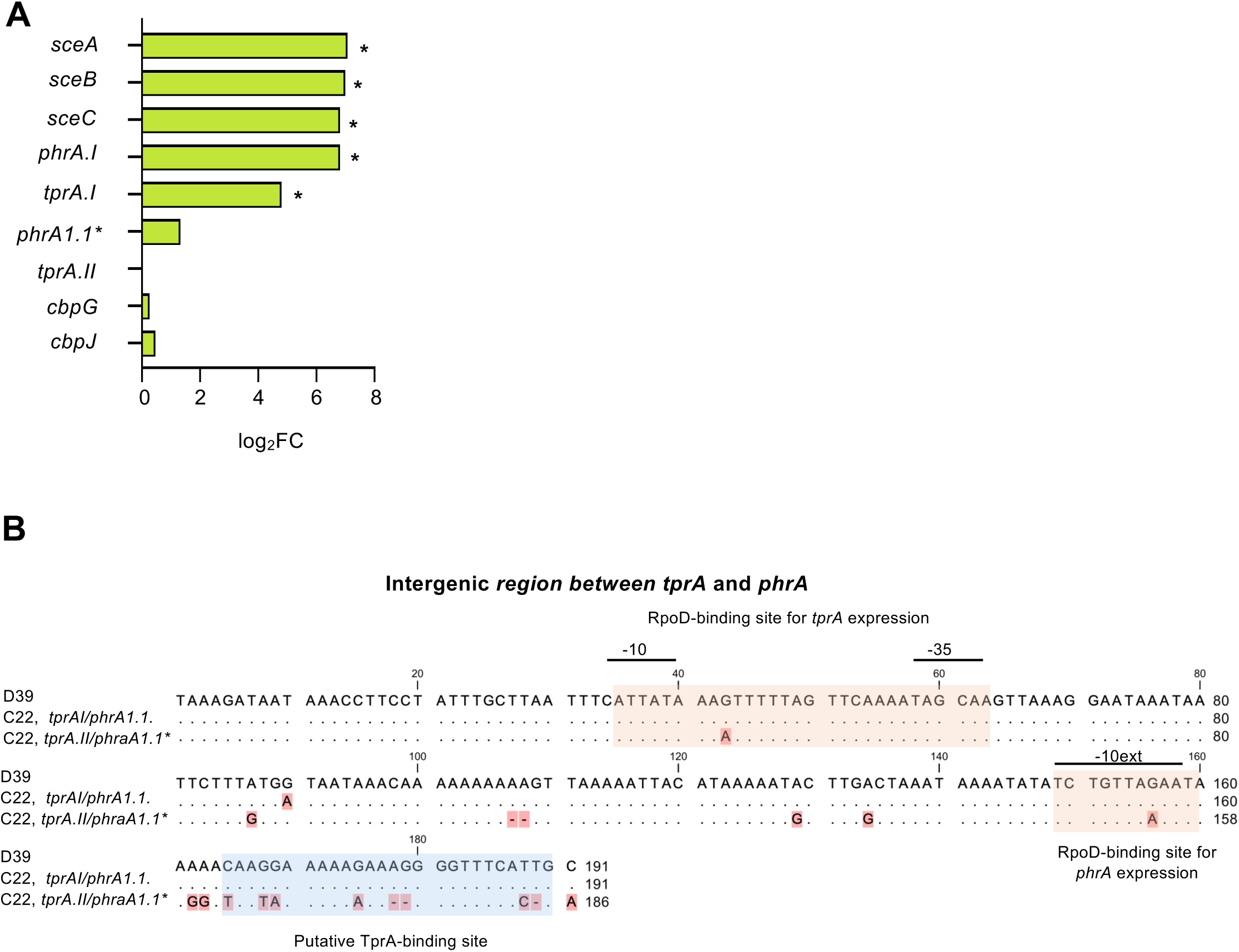
PhrA1.1 activates gene expression of *sce* locus, including *tprA.I*, but not of its paralogue *tprA.II*. Evaluation of PhrA1.1 role in control of gene expression in *S. mitis* strain C22WT was done by RNA-seq. C22WT was grown in CDMgal, with and without synthetic PhrA1.1. Following incubation, RNA was extracted. **A.** Transcriptomic analysis of *tprA* paralogues and their downstream genes after PhrA stimulation. Differential expression was observed only for genes located downstream of the *tprA.I* (bars with *). In contrast, expression of its paralogue, *tprA.II,* and its associated genes, *cbpG/J* remained unchanged. Cut-off thresholds for significance: FDR < 0.05, absolute log₂fold change > 3, and P-value < 0.001. The global transcriptomic profile of strain C22 plus PhrA1.1 can be found in Table S4. **B.** Alignment of the intergenic regions between *tprA-phrA* paralogs in strain C22 and strain D39 from *S. pneumoniae*. Key regulatory elements and binding sites for transcriptional initiation are annotated. Nucleotides shown as dots are the same as the reference sequence (D39). Nucleotides with a red background indicate differences from the reference. Dashes represent alignment gaps.

Together these results show that the two copies of TprA in *S. mitis* C22 are functional, and both repress the *sce* locus with additive effects. In addition, PhrA can release inhibition for both paralogues. This pheromone, however, induces expression of *tprA.I* but not of *tprA.II*.

### *In silico* analyses suggest that differences in intergenic sequences underlie distinct regulation of *tprA.I* and *tprA.II*

To investigate why the pheromone PhrA induced expression of TprA.I but not of TpA.II, we compared the intergenic sequences upstream of both *tprA* paralogues in strain C22 with that of strain D39, where *tprA/phrA* expression is active and peptide-dependent (12). The intergenic region of *tprA.I/phrA1.1* closely resembled that of D39, whereas the *tprA.II/phrA1.1\** region differed significantly, particularly in the palindromic sequence predicted to be the TprA binding site (**Fig. 7B**). These findings suggest that differences in the intergenic regions, especially in the palindromic binding site, are driving the differential gene regulation observed for the two copies.

### The two TprA paralogues act in concert to repress bacteriocin production and competence induction, coordinating stress and metabolic homeostasis

To explore the regulons of the two TprA paralogues, we performed RNA-seq on C22*ΔtprA.I,* C22*ΔtprA.II*, and the double mutant C22*ΔtprA.I/ΔtprA.II*, and compared its transcriptomes to that of the parental strain C22WT (**Table 1, Table S6**). In total, significant changes in gene expression were detected for 183 genes across one or more mutants. Among the various expression patterns identified, the most frequent ones corresponded to redundant repression by TprA.I and TprA.II (where upregulation of a gene occurred only in the double mutant C22*ΔtprA.I/ΔtprA.II*; observed in 44 genes), independent repression (in which gene expression increased in all mutants; 33 genes), and paradoxical epistasis (where single deletions of *tprA.I* or *tprA.II* increased expression of a given gene but the double mutant restored wild-type levels). Additionally, some results revealed that both TprA.I and TprA.II can also act as activators in which case, deletion of one or the other resulted in lower gene expression than in the C22WT.

**Table 1.**
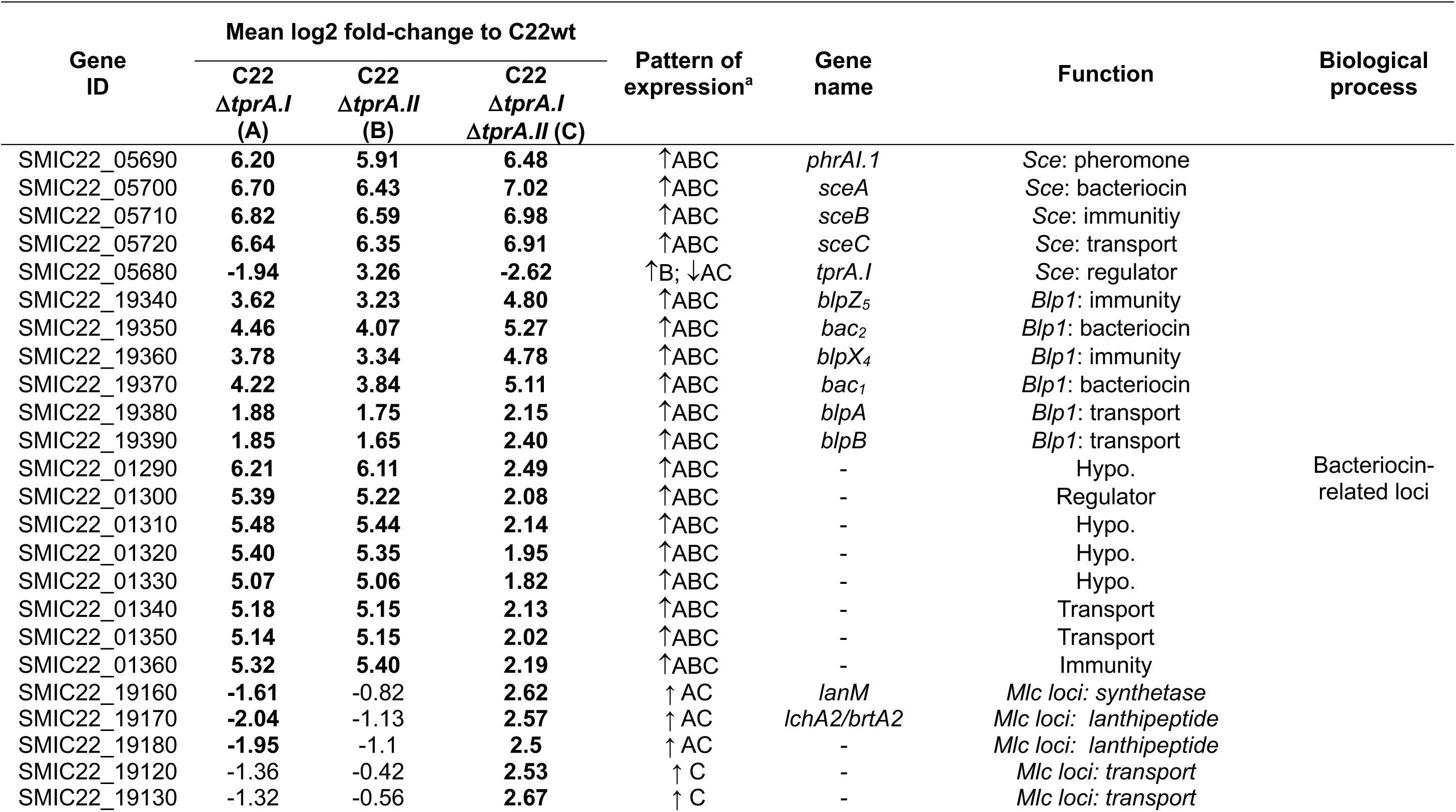
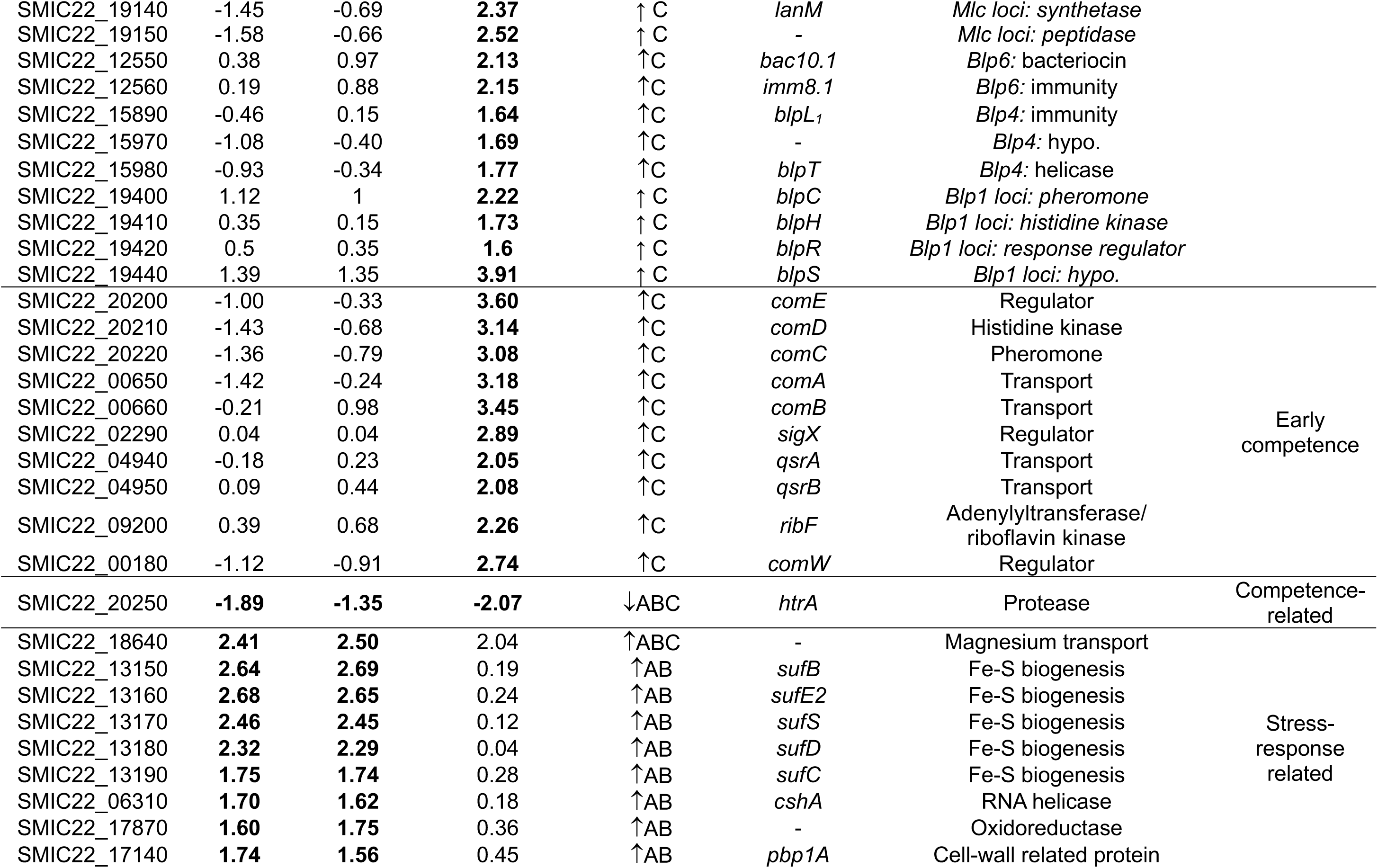
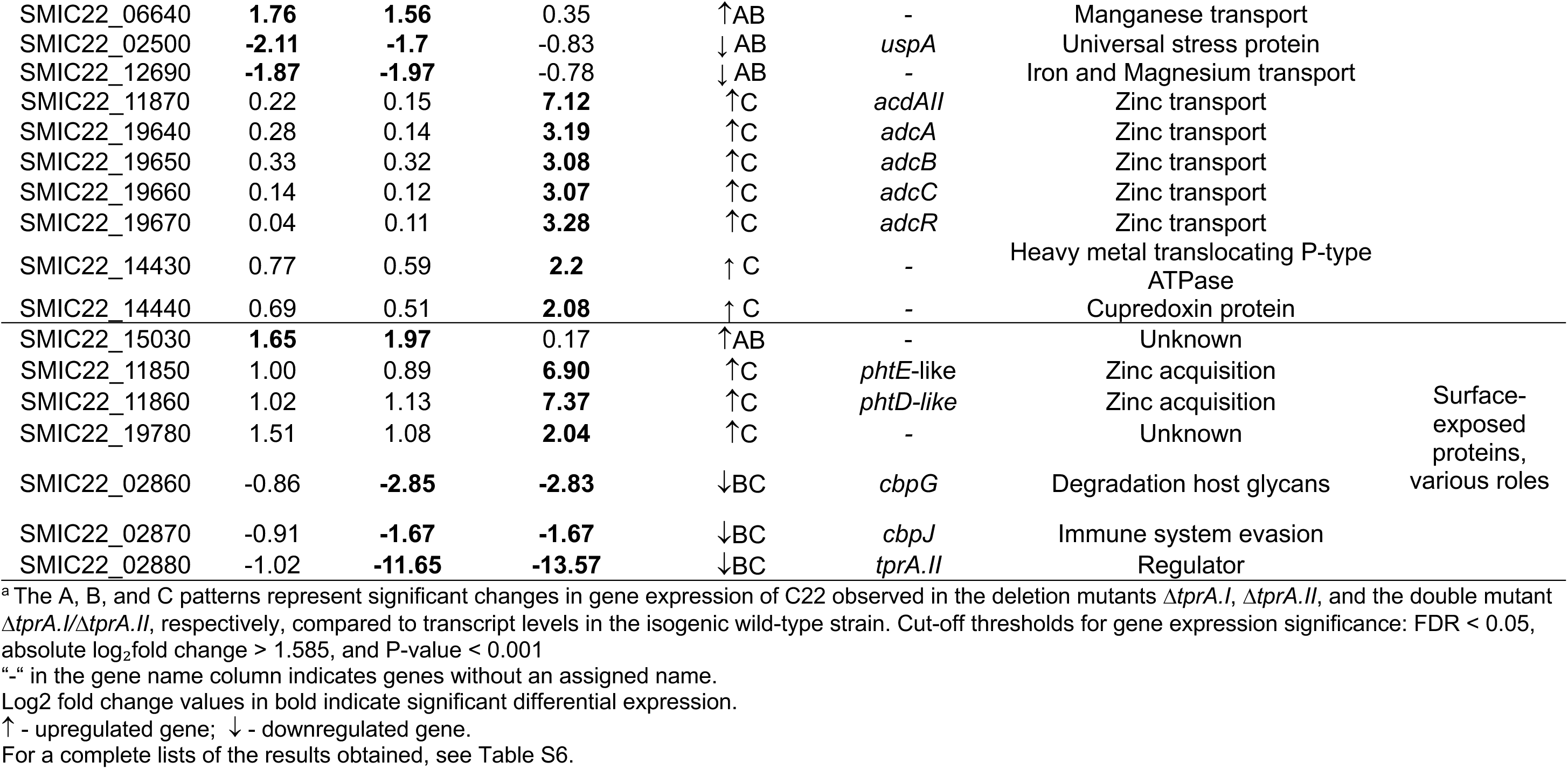
Differentially expressed genes in strains C22*11tprA.I*, C22*11tprA.II*, C22*11tprA.I11tprA.II* associated with bactericidal activity, competence, stress response, and other functions.

RNA-seq analysis revealed that deletion of *tprA.I* or *tprA.II* resulted in a 64-fold increase in expression of the streptococcin E (*sce*) locus, located downstream of *tprA.I*. The double *tprA.I/tprA.II* mutant showed a further increase of c.a. 128-fold. These results demonstrated that the two TprA paralogues act as additive repressors of the *sce* promoter. In contrast, the results indicated that expression of *cbpG* and *cbpJ,* located downstream of *tprA.II*, was modestly activated by TprA.II (3-fold) and not affected by TprA.I. These observations are in line with the observed differences in the intergenic regions of the two systems (see above, **Fig. 7B**).

Deletion of both *tprA* paralogues, but not the single deletions, promoted early competence induction (**Table 1**). Compared to the wild-type, the double mutant showed increased transcription of the genes encoding for the competence stimulating peptide (CSP) and its associated two-component regulatory system (*comCDE*), the CSP transporter (*comAB*), and the alternative sigma factor driving the competence regulon (*sigX*). We also detected downregulation of the *htrA* protease that degrades CSP (29), consistent with sustained activation of competence in the double mutant. Other competence-associated effectors were also upregulated, including bacteriocin related genes (*blp4* and *mlc*) and *qsrA/B* and *ribF*, which contain a ComE binding site in their promoters (30). Additionally, strong induction was detected for the two major zinc transport operons, the *adcABCR* and *adcAII* (31, 32). In contrast, expression of a polysaccharide lyase gene, potentially involved in host glycan degradation and colonization fitness (33), was markedly reduced. Collectively, these results indicate that TprA.I and TprA.II act as redundant repressors of competence-related genes.

Moreover, we identified a subset of 28 genes that were upregulated and 19 genes that were downregulated in either *ΔtprA.I* or *ΔtprA.II*, but not in the double mutant *ΔtprA.I/ ΔtprA.II* (**Table 1, Table S6**). This set included several genes involved in stress response, such as the *sufCDE2B* operon, a DEAD-box RNA helicase, manganese transport genes, and a LLM flavin-dependent oxidoreductase (34–37). The specific expression changes observed in the single mutants, but not in the double mutant, suggest an indirect effect of the TprA paralogues. This may result from partial or premature competence induction (described above), which transiently alters cellular metabolism and redox balance, leading to a competence-associated secondary stress response. In the double mutant, where competence is fully and synchronously induced, this secondary response appears to be bypassed, normalizing oxidative stress-related gene expression.

As expected, RNA-seq also revealed a subset of genes, 32 upregulated genes and 12 downregulated, that were differentially expressed in both the single and double mutants (*ΔtprA.I, ΔtprA.II, ΔtprA.I/ ΔtprA.II)* (**Table 1, Table S6**). Genes consistently upregulated in all mutants included those involved in bactericidal functions (the *sce* locus discussed above and the *blp1* locus), and also an uncharacterized locus predicted to encode short peptides and immunity proteins, indicating that both TprA paralogues act as repressors of these loci.

## DISCUSSION

The TprA/PhrA signaling system has emerged as a key regulatory circuit linking bacterial communication with metabolic control and competitive behavior in *Streptococcus pneumoniae* (12–14). Here, we show that orthologues of this system are widely distributed among mitis group streptococci and that it has been both conserved and diversified in *S. mitis*. Using *S. mitis* strain C22 as a model, our results reveal that this commensal species can harbor two functional TprA paralogues, providing additional regulatory flexibility to fine-tune social behavior such as competence induction, bacteriocin production, and stress response. Moreover, our findings expand the known TprA/PhrA regulon to include loci involved in adhesion, antibiotic resistance, and ABC transporters, implying a broader role for this system in shaping microbial fitness and interactions within polymicrobial communities of the upper respiratory tract.

Our comparative genomic analyses show that the TprA/PhrA system is largely confined to members of the mitis group, consistent with its previously proposed role in adaptation to the upper respiratory tract (13, 38). Across the *S. mitis* pangenome, a substantial fraction of genomes encode *tprA* but lack *phrA*, suggesting that some strains may rely on heterologous PhrA signals from co-colonizing bacteria or maintain PhrA-independent regulation. This diversity supports the idea that quorum sensing in *S. mitis* involves both self and non self-communication, enabling coordination of behavior within polymicrobial communities (6, 39, 40).

In both *S. pneumoniae* and *S. mitis*, we found that the genomic region downstream of *tprA* is highly variable and frequently includes bacteriocin-related loci such as the streptococcin E (*sce*) cluster, which encodes a lactococcin 972–like bacteriocin (26, 41). While we show that the *sce* locus confers a measurable fitness advantage in strain competition assays, our genomic data also uncovered the frequent occurrence of incomplete *sce* loci, typically lacking the *sceA* bacteriocin gene but retaining the *sceB* and *sceC* immunity and transport genes. These incomplete configurations suggest the existence of “cheater” phenotypes (6, 42), in which strains avoid the metabolic burden of bacteriocin production yet remain protected by expressing the corresponding immunity proteins. The widespread retention of *sceBC* across *S. mitis* genomes further implies a network of cross-protection among related bacteriocins, facilitating coexistence among closely related strains within polymicrobial communities (26, 42). Such asymmetric distribution of bacteriocin and immunity genes, regulated by TprA, likely contributes to maintaining microbial diversity and stability in the oral and nasopharyngeal ecosystems. Indeed, these niches are known to harbor remarkable species- and strain-level diversity of streptococci (38, 43, 44).

Functionally, we further show that the *sce* locus in *S. mitis* C22 encodes a PhrA-inducible bacteriocin that confers a competitive advantage both *in vitro*, in biofilm competition assays, and *in vivo*, in a *Galleria mellonella* infection model. These findings extend the role of the TprA/PhrA system beyond *S. pneumoniae*, where previous work showed that TprA controls lantibiotic synthesis and peptide-mediated signaling (12), adding evidence that this quorum-sensing circuit also governs bacteriocin-mediated competition (streptococcin E) and social interactions in commensal streptococci.

The discovery that some strains contain two *tprA* copies (here described using *S. mitis* C22), both functional and responsive to the same pheromone, reveals a new layer of regulatory complexity not described before. Our transcriptomic data indicate that the TprA paralogues display partially overlapping but distinct regulons, with one (*tprA.I*) responding to PhrA-dependent activation while the other (*tprA.II*) remains largely unresponsive to PhrA but exerts constitutive repression. This unidirectional network, where TprA.II inhibits *tprA.I* but is not reciprocally repressed, mirrors the asymmetric regulation described for the TprA2/PhrA2 system of *S. pneumoniae* PMEN1, suggesting a conserved evolutionary logic in the emergence of layered quorum-sensing circuits (10). Both paralogues repress the *sce* locus in an additive manner and control competence development, linking bacteriocin regulation with horizontal gene transfer and bacterial evolution. The co-repression of these traits suggests that the TprA paralogues act as a checkpoint system coordinating metabolic investment and stress response, ensuring that energy-demanding processes such as bacteriocin production and competence are activated only under specific community or environmental cues.

Differences in the intergenic regions of the two *tprA/phrA* copies in C22 appear to underlie their divergent responsiveness to PhrA. Whereas the upstream region of *tprA.I* resembles that of *S. pneumoniae* D39 (12), enabling peptide-dependent activation, *tprA.II* shows alterations in the palindromic TprA-binding site, likely decoupling it from pheromone feedback. These sequence variations may represent evolutionary fine-tuning, allowing strains with both paralogues to balance redundancy and specialization in quorum-sensing regulation. This regulatory asymmetry may also explain the differential enrichment of stress-related genes across *S. mitis* C22 mutants: both single *tprA* deletions upregulated oxidative stress response pathways, while the double mutant displayed enhanced *ribF* expression, potentially linking competence activation with redox homeostasis through FAD metabolism (45, 46).

Our findings highlight the potential for interspecies communication between *S. mitis* and *S. pneumoniae*. Analysis of *phrA* allelic diversity revealed both shared and species-specific variants, with some alleles (e.g., *phrA1.1*) enabling cross-species activation and others maintaining species-restricted signaling. Consistent with this, exogenous PhrA peptides from *S. mitis* activated TprA signaling in *S. pneumoniae*, and vice versa, demonstrating that diffusible pheromones mediate reciprocal sensing between species. Likewise, *S. mitis* supernatant triggered *tprA/phrA* expression in *S. pneumoniae*, confirming that these peptides act as interspecies cues. Such interspecies signaling could influence population structure, virulence activation, or even colonization hierarchy, as *S. mitis* has been reported to modulate pneumococcal behavior and the growth of other respiratory pathogens through metabolic and oxidative interactions (6, 47). These findings suggest that, in mixed communities, commensal *S. mitis* may modulate pneumococcal behavior, influencing colonization dynamics, horizontal gene transfer, and transitions to pathogenic states. Conversely, allele-specific signaling barriers likely limit interference among co-colonizing strains, preserving species identity within complex oral and nasopharyngeal biofilms.

Collectively, our data support a model in which TprA/PhrA systems act as conserved communication hubs coordinating competition, competence, and stress adaptation across the mitis group (see model in **Fig. 8**). In *S. mitis*, the expansion of this circuitry through acquisition and diversification of homologous loci provides a molecular basis for nuanced regulation and interspecies dialogue with *S. pneumoniae*. We propose that such communication contributes to the structuring of microbial communities in the upper respiratory tract and may indirectly influence pneumococcal virulence and persistence. This work redefines the TprA/PhrA regulon as a central integrator of bacterial social behavior and highlights pheromone-mediated crosstalk as a key driver of community structure and evolutionary innovation within the human respiratory microbiota.

**Figure 8.**
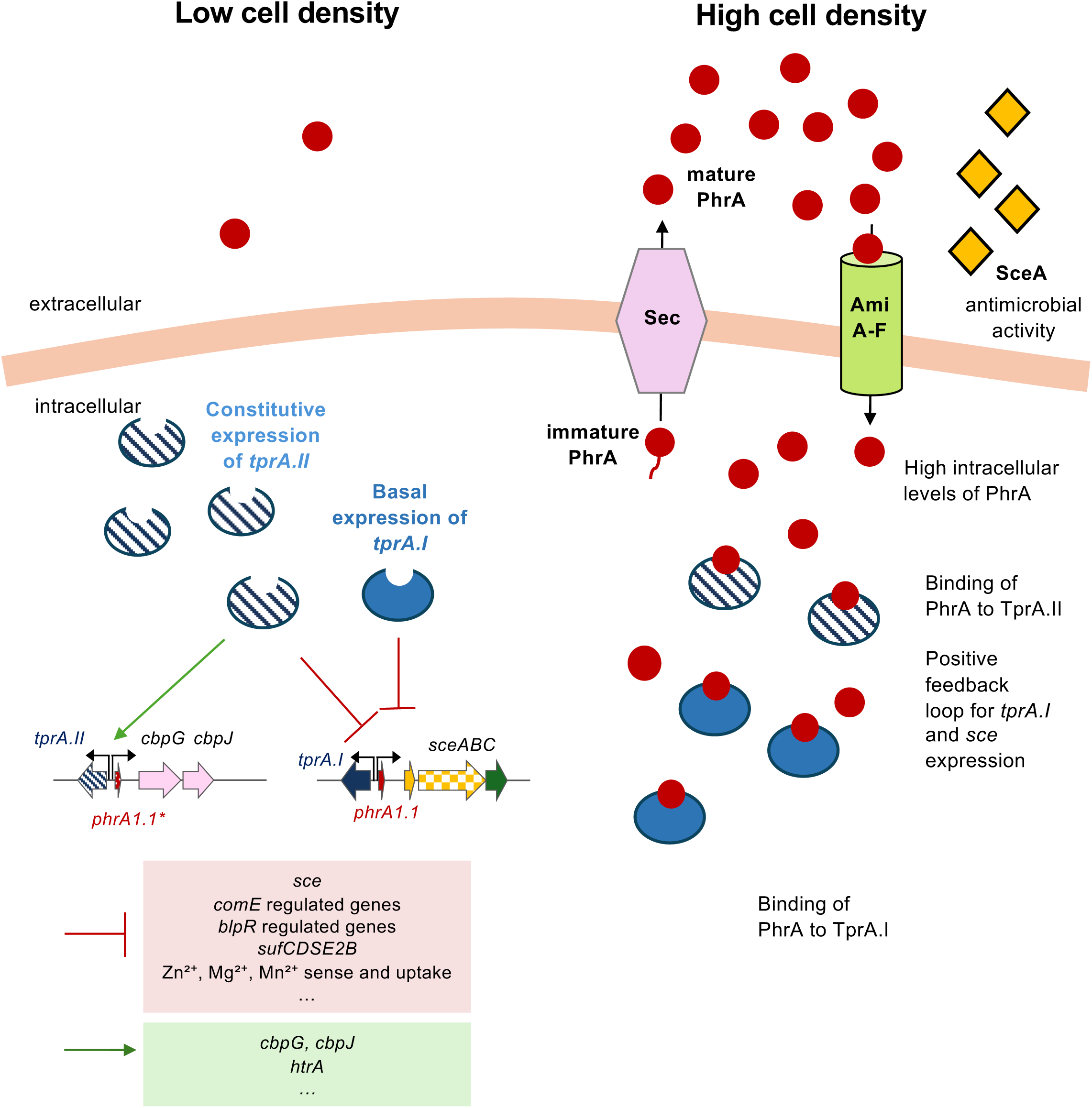
Model of TprA paralog-mediated gene regulation in *S. mitis* C22WT. This model depicts a dual regulatory network governed by the *tprA* paralogs, TprA.I and TprA.II, which act as transcriptional repressors and modulate distinct yet overlapping regulons in response to cell density during nasopharyngeal colonization. At low cell density, when quorum-sensing systems are inactive, TprA.II is constitutively expressed and represses *tprA.I*, *phrA.I*, and the *sceABC* operon, while activating expression of downstream choline-binding protein genes (*cbpG*, *cbpJ*) potentially involved in adhesion. TprA.II does not repress its own expression. In contrast, TprA.I represses its own operon (*tprA.I–phrA.I*) and the adjacent *sceABC* locus, establishing an autoregulatory loop that maintains low expression under non-inducing conditions. As bacterial density increases, the signaling peptide PhrA is secreted via the Sec pathway, processed extracellularly, and re-imported through the AmiA–F transporter. Accumulation of intracellular mature PhrA interferes with TprA.II-mediated repression, leading to derepression of *tprA.I* and *sceABC*. Activated *tprA.I* then reinforces its own regulatory feedback, fine-tuning expression levels of its regulon. This multilayered regulatory system coordinates population-dependent control of bacteriocin production, competence, and stress adaptation. Increased expression of *sceABC* results in production of the antimicrobial peptide SceA, which confers a competitive advantage in mixed biofilms and *Galleria mellonella* infection models. The distribution of *sce* immunity genes among *S. mitis* and *S. pneumoniae* suggests selective protection within these species, supporting a role for this system in microbial competition and persistence in the nasopharyngeal niche.

## ACKNOWLEDGMENTS

This work was supported by FCT - Fundação para a Ciência e Tecnologia, I.P., through project STOPneumo (PTDC/BIA-MIC/30703/2017), MOSTMICRO-ITQB R&D Unit (UIDB/04612/2020, UIDP/04612/2020), LS4FUTURE Associated Laboratory (LA/P/0087/2020) and by european funds from FEDER - “Fundo Europeu de Desenvolvimento Regional”. B. Ferreira was supported by a PhD fellowships (2020.05293.BD) from FCT.

We thank Adriano Henriques (ITQB NOVA) and Cristina Silva Pereira (ITQB NOVA) for interesting and productive discussions, and Karina Mueller-Brown for her assistance with the peptide assays. Finally, we would like to thank Melissa Jansen van Rensburg and Angela Brueggemann for advise on the selection of curated PubMLST sequences for *S. pneumoniae*.

## CONTRIBUTIONS

RSL, CV and LH were responsible for the concept and design of the study. RSL contributed with study collections, reagents and materials. LH and HY contributed with reagents and materials. BF performed all experimental work except for competition assays in *Galleria mellonela* infection model, which were performed by OG. SC contributed to comparative genomics analysis. Data analysis and interpretation of results was done by BF, CV, LH and RSL. The manuscript was drafted by BF, CV, LH and RSL and critically revised by all authors. All authors read and approved the final version of the manuscript.

---

**Figure S1.**
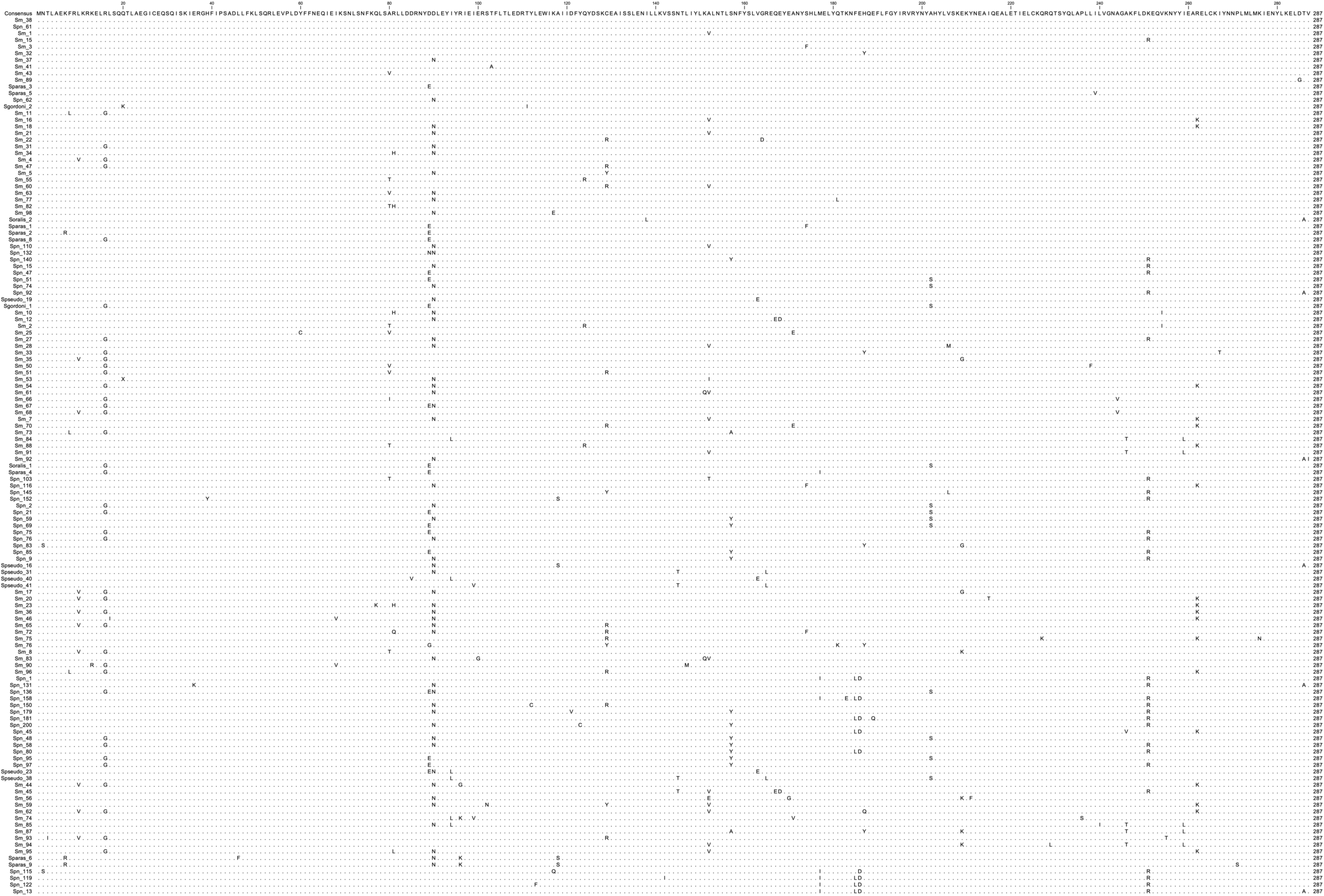
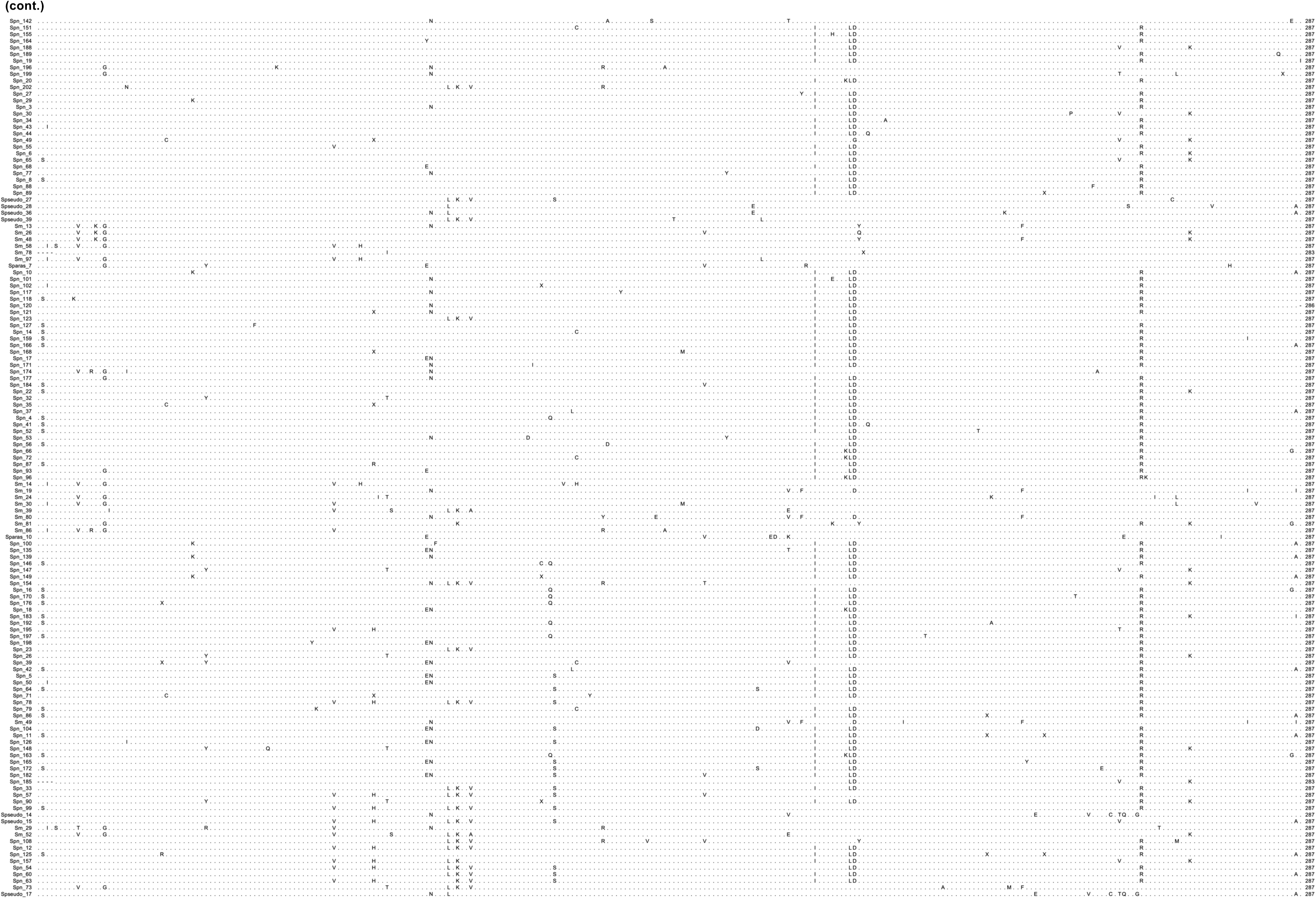
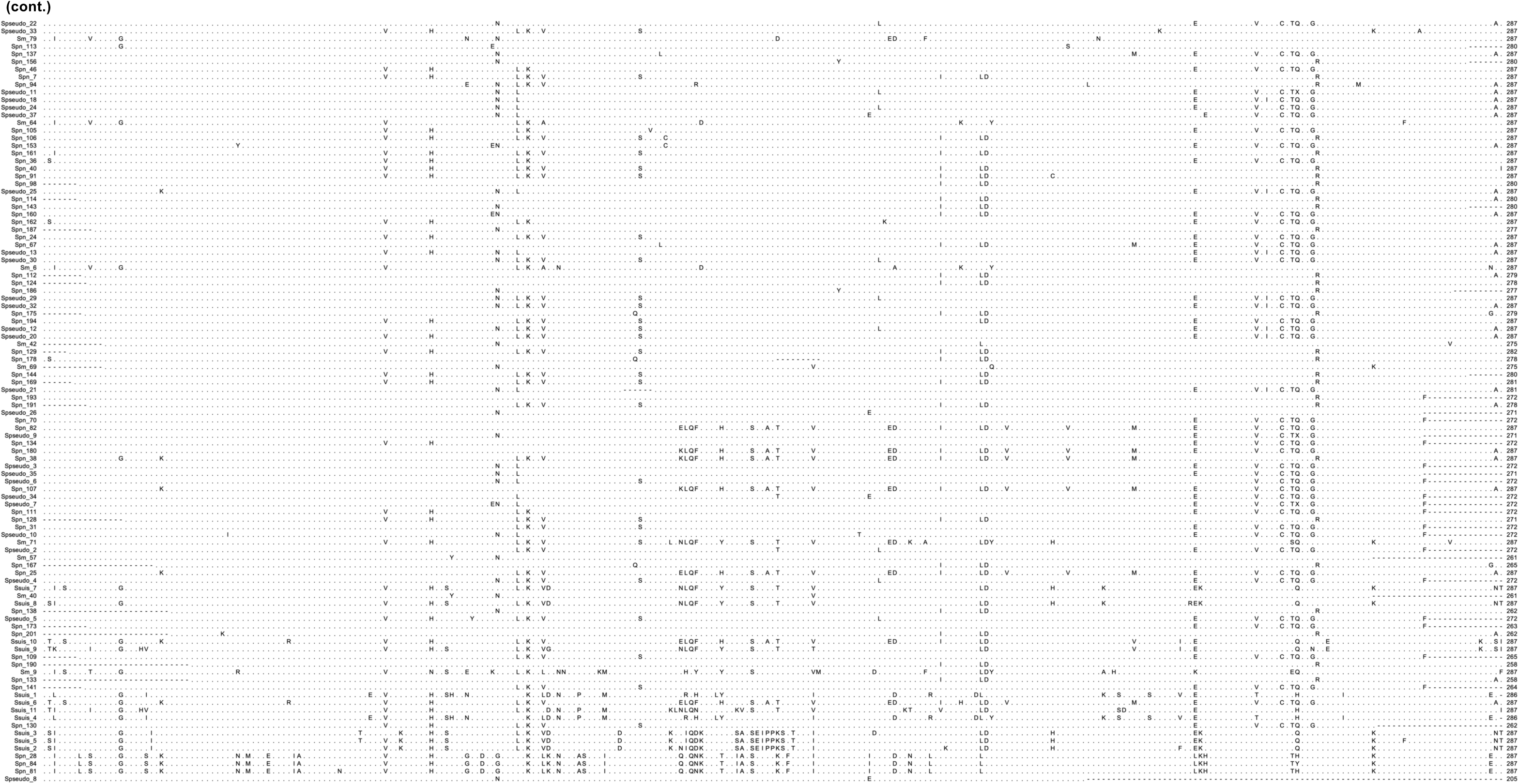
Alignment of TprA variants across oral streptococci. TprA variants were defined at single-amino-acid resolution, where sequences differing by ≥1 residue were treated as distinct, and identical sequences were collapsed to one representative. The resulting dataset comprised unique TprA sequences from *Streptococcus pneumoniae* (n=202), *S. mitis* (n=98), *S. pseudopneumoniae* (n=41), *S. oralis* (n=2), *S. parasanguinis* (n=10), and *S. suis* (n=11). A global consensus across all variants (“Consensus”) was obtained and used as the alignment reference. In total, 366 variants were aligned, six of which are shared across species (Spseudo_1–Spn_70; Sgordoni_1–Soralis_1–Spn_21; Spn_61–Sm_38; Spn_62–Sm_37; Spn_76–Sm_27; Spn_110–Sm_21). Variant prefixes denote species: Spn_ (*S. pneumoniae*), Sm_ (*S. mitis*), Spseudo_ (*S. pseudopneumoniae*), Soralis_ (*S. oralis*), Sparas_ (*S. parasanguinis*), Sgordoni_ (*S. gordonii*), Ssuis_ (*S. suis*).

**Figure S2.**
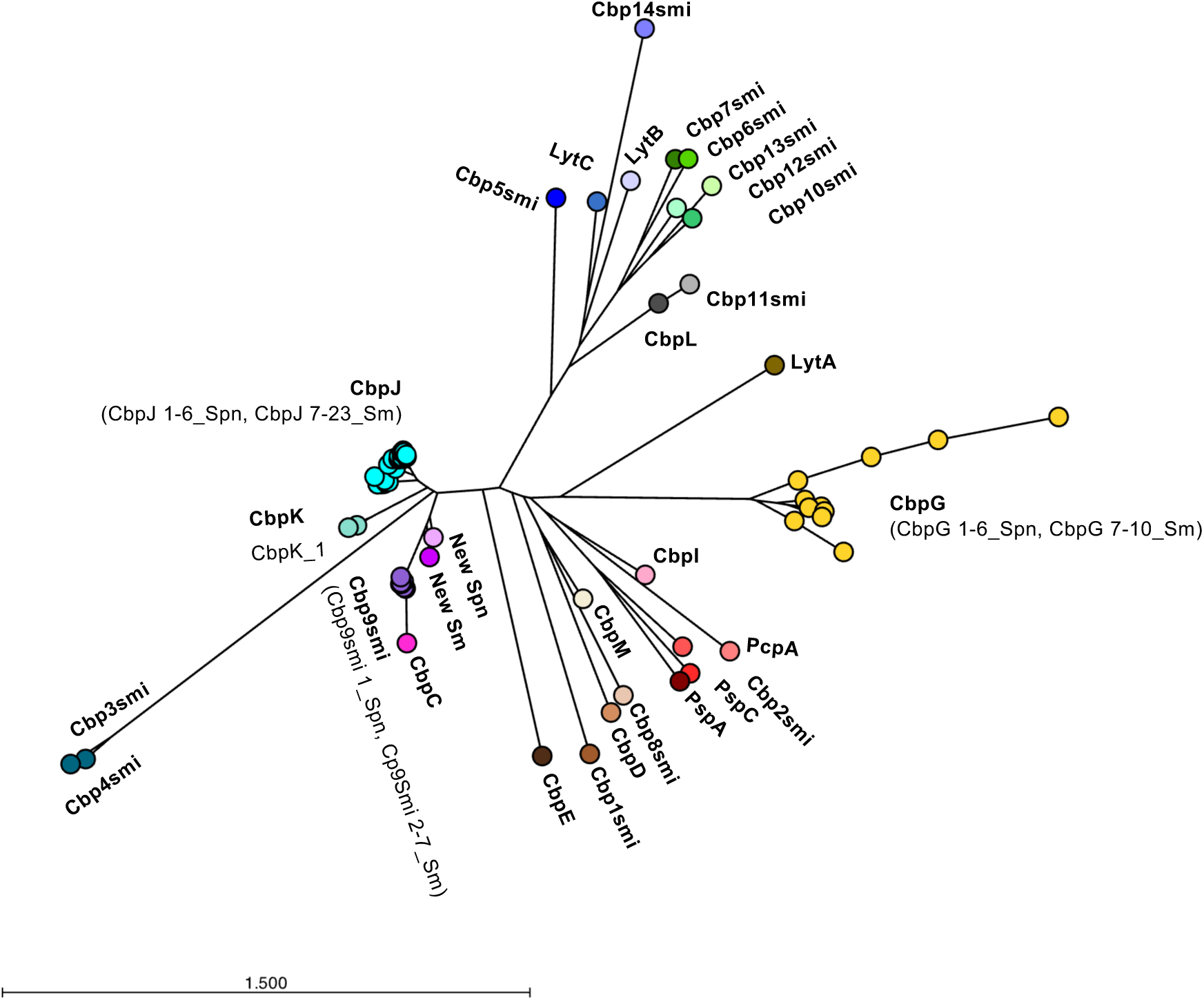
Phylogenetic tree of choline binding proteins (Cbp) located downstream of *tprA* in *S. pneumoniae* and *S. mitis*. A neighbor-joining tree was constructed using the Jukes-Cantor model, including Cbp proteins found downstream of *tprA* in *S. pneumoniae* and *S. mitis* as well as reference Cbp from both species available in the NCBI database. The scale bar represents the number of amino acid substitutions per site.

**Figure S3.**
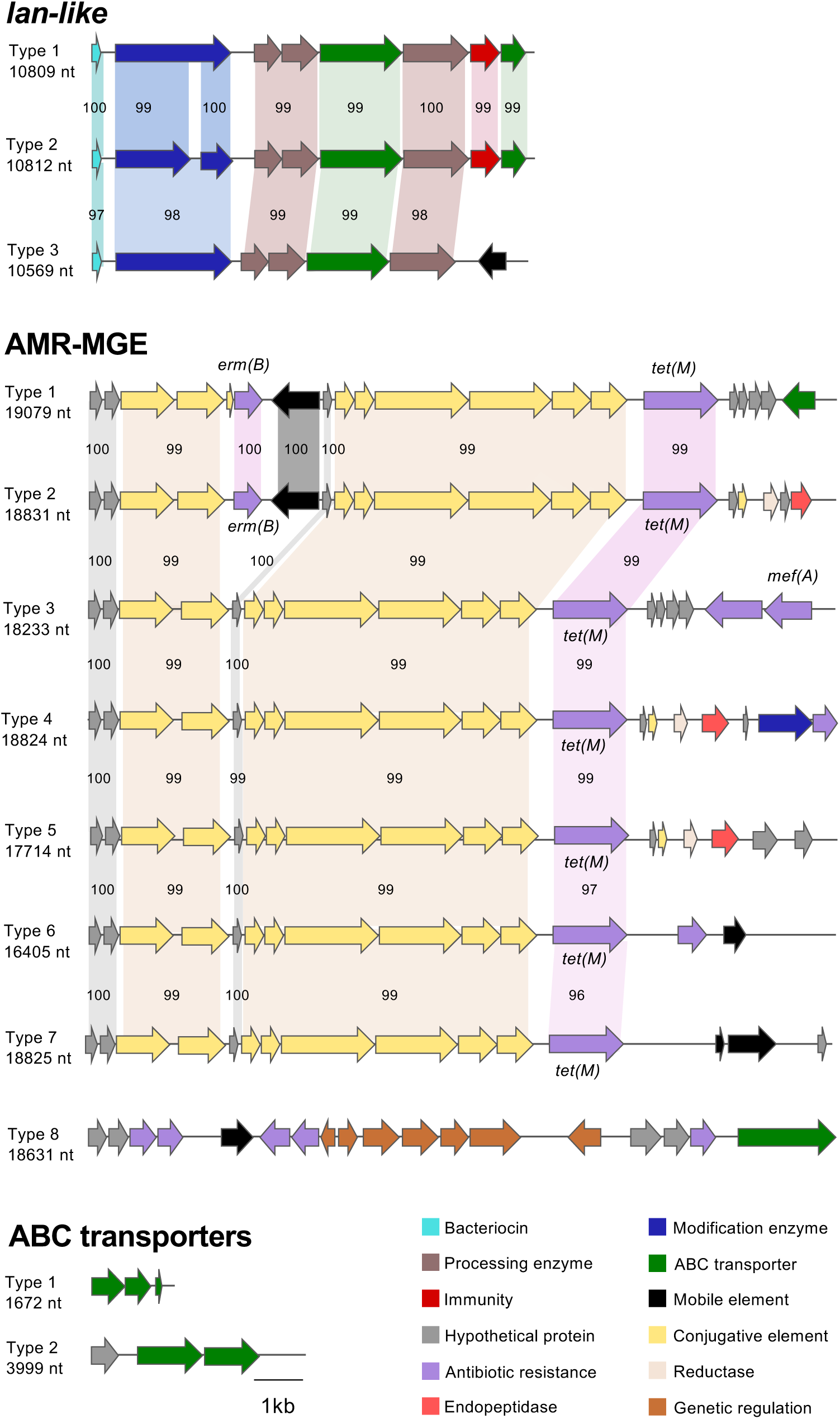
Genetic organization of the *lan-like*, AMR-MGE, and ABC-transporter loci located downstream of *tprA* in *S. pneumoniae* and *S. mitis*. Results obtained based on analysis of 7,548 *S. pneumoniae* and 322 *S. mitis* genomes. For comparison, type-1 locus is shown as the reference for lan-like and AMR-MGE regions. Geometric forms spanning across loci indicate segments conserved between different locus types. Gene function is indicated by color, and the lateral sizes (nt) correspond to the total represented regions. Percent values indicate nucleotide identity relative to the type-1 reference locus. *Lan-like*, lantibiotic-like region; *AMR-MGE*, antimicrobial-resistance mobile genetic element.

**Figure S4.**
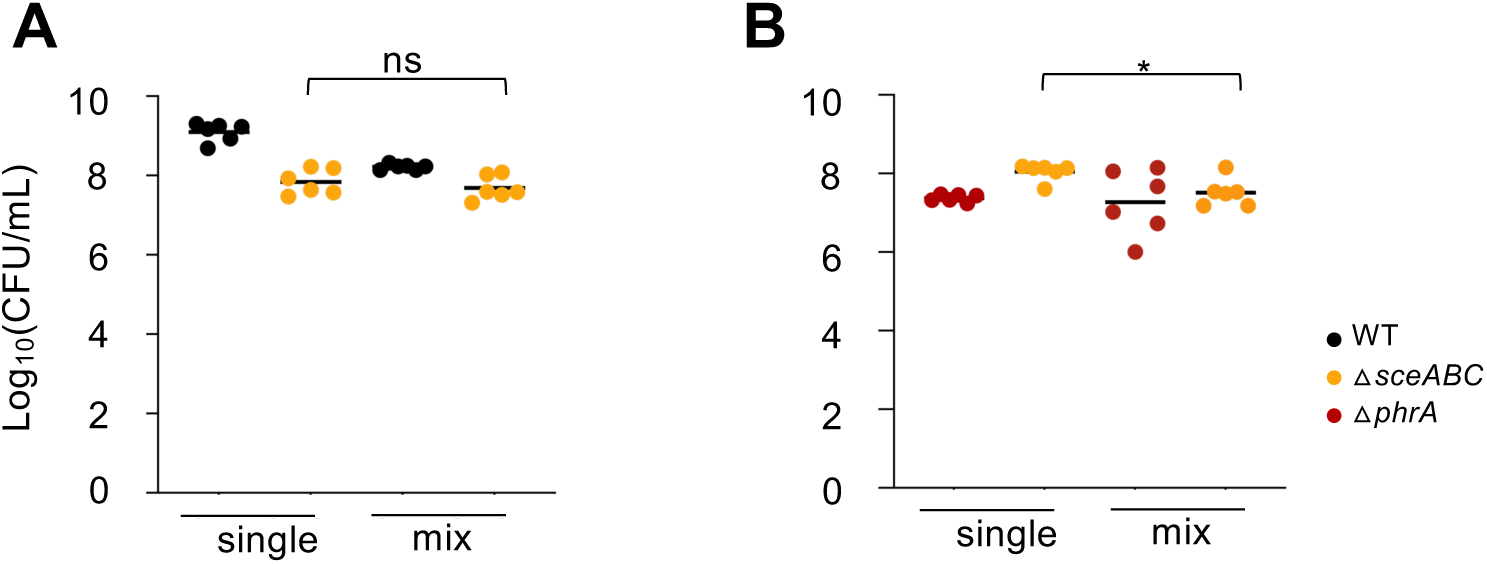
The *sce* locus exhibits antimicrobial activity, with PhrA regulating its inhibitory potential. **A.** The antimicrobial activity of the *sce* locus was assessed in an *in vitro* biofilm model by co-culturing C22WT (WT) and its isogenic △*sceABC* mutant at a 1:1 ratio (initial inoculum of 10^5^ CFU/mL) in CDM galactose. At 24h, no significant effect was detected. **B.** The role of PhrA on *sce* locus inhibition was evaluated using strain C22Δ*phrA (*Δ*phrA)* instead of WT. Bacterial loads quantification was done at 24h by plating the initial inoculum and the final biofilm on blood agar (total load) and blood agar supplemented with kanamycin (selective for △*sceABC* mutant). P-values were calculated by paired Student’s t-test. Strains tested are color-coded: black – WT; yellow - △*sceABC*; red - △*phrA.* \**P-value* <0.05, ns, not significant. All assays were performed at least in triplicates, with graphics showing mean ± SEM.

**Figure S5.**
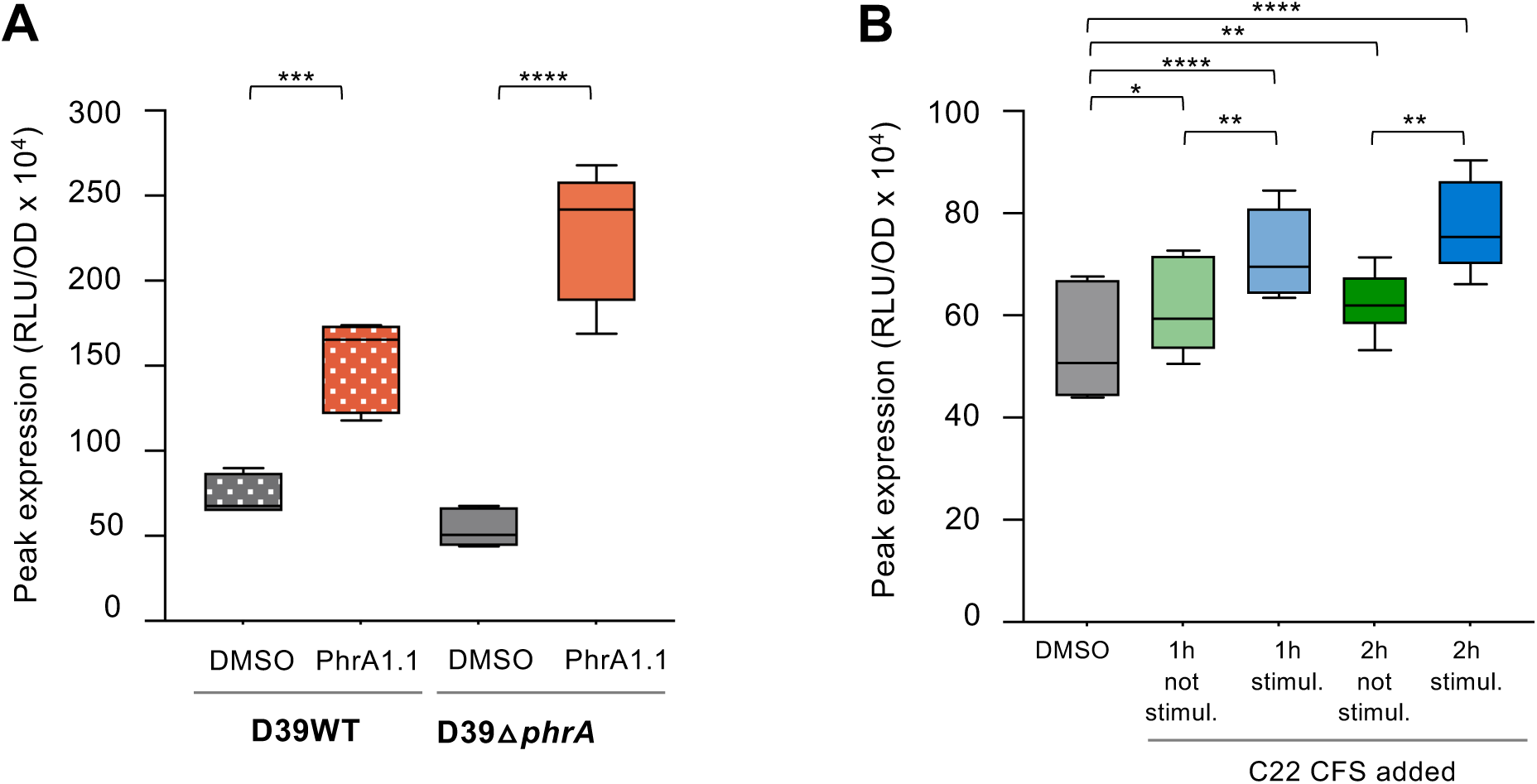
Expanded analysis of inter-species communication via the *tprA/phrA* system. Activation of the *phrA* promoter in *S. pneumoniae* D39 background strains, each carrying a PphrA:mNeonGreen-luciferase fusion at the *cil* locus. **A.** Reporter activation following direct addition of 1 mM synthetic PhrA1.1 C10 peptide or DMSO to cultures of D39WT or D39Δ*phrA*. Strains tested and conditions are color-coded: dotted grey, D39WT+DMSO; dotted orange, D39WT+PhrA1.1; full grey, D39Δ*phrA*+DMSO; full orange, D39WT+PhrA1.1. **B.** Reporter activation of D39△*phrA* in response to cell-free supernatants (CFS) from *S. mitis* C22WT, either stimulated with 1 mM synthetic PhrA1.1 C10 or untreated (with DMSO). Following stimulation, cultures were washed to remove residual peptide and regrown for 1 or 2 hours before collecting and filtering the supernatants. D39Δ*phrA* reporter strain was incubated with the resulting CFS or DMSO (as a control). Assays were conducted in 96-well plates at 37 °C with 1 s orbital shaking. OD_595_ and luminescence (RLU/OD) were measured every 10 minutes over a 16-hour period using a BioTek Neo2 plate reader. Conditions tested are color-coded: grey, DMSO; light green, 1h CFS from non-stimulated C22WT; light blue, 1h CFS from stimulated C22WT; intermediate green, 2h CFS from from non-stimulated C22WT; dark blue, 2h CFS from stimulated C22WT. Data represent the mean ± SEM from three biological replicates. P-values were calculated by two-way ANOVA. \**P-value* <0.05, \*\**P-value* <0.01, \*\*\**P-value* <0.001; \*\*\*\**P-value* <0.0001.

